# Mechanism for oil-phase separation by the lipid droplet assembly complex

**DOI:** 10.1101/2025.08.12.669882

**Authors:** Pedro C. Malia, Siyoung Kim, Yohannes Ambaw, Gregory A. Voth, Tobias C. Walther, Robert V. Farese

**Affiliations:** Cell Biology Program, Sloan Kettering Institute, Memorial Sloan Kettering Cancer Center, New York, NY, USA; Howard Hughes Medical Institute, NY, USA; Pritzker School of Molecular Engineering, The University of Chicago, Chicago, IL, USA; Department of Chemistry, Chicago Center for Theoretical Chemistry, James Franck Institute, and Institute for Biophysical Dynamics, The University of Chicago, Chicago, IL, USA

**Author notes:** These authors contributed equally (Tobias C. Walther, Robert V. Farese). **Contact information:** Corresponding authors: Tobias C. Walther and Robert V. Farese Jr.

## Abstract

Cells store metabolic energy as triglyceride (TG) oils in lipid droplets (LDs). LDs form *de novo* from the endoplasmic reticulum. How the lipid droplet assembly complex (LDAC), composed of seipin and LDAF1^1,2^, catalyzes the organized formation of an oil phase in a membrane bilayer before spontaneous phase separation is triggered is unknown. Here, we reconstitute LD formation *in vitro* using purified LDAC and membranes containing physiologic levels of TG, demonstrating that the LDAC is both necessary and sufficient to catalyze oil-phase formation below the threshold of spontaneous phase separation. Structural studies of the LDAC reveal that LDAF1 forms a central ring within a seipin cage, creating a toroidal, membrane-spanning structure. Molecular dynamics simulations and biochemical assays show that this structure forms a selective chamber within the ER bilayer that limits phospholipids but allows TG to access a reaction compartment between the inner and outer rings of the LDAC. Within this compartment, TG interacts with LDAF1 and each other to form an oil phase to initiate LD formation. Thus, the LDAC acts as a protein catalyst for oil-phase separation in cells, revealing a fundamental mechanism for how cells resolve the biophysical challenge of storing oils within a hydrophilic environment in an organized manner.

---

Most organisms store metabolic energy as reduced carbons in an organic oil composed primarily of triacylglycerols (TGs). TGs and other neutral lipids^3^ are synthesized in the endoplasmic reticulum (ER) and released into the membrane bilayer^4–7^. TGs are thought to accumulate within a membrane bilayer till they reach a critical concentration (i.e., ∼3 mol% of membrane lipids, as measured by nuclear magnetic resonance spectroscopy^8^) at which point they phase-separate to form an oil lens^9^.

Cells have evolved dedicated protein machinery to facilitate lipid droplet (LD) formation. Such control is crucial to preserve ER function and cellular homeostasis. In particular, the fidelity of LD formation maintains ER membrane integrity and supports its functions, such as calcium homeostasis^10,11^. Disruption of normal LD formation activates stress responses^12^ and, in susceptible cells such as adipocytes, triggers dysfunction and cell death^13^. Thus, regulated LD formation is an essential evolutionary adaptation to manage neutral lipid synthesis and storage.

The core machinery that mediates LD formation is the LD assembly complex (LDAC), composed of the oligomeric proteins seipin and its binding partner LDAF1^1,2,14^. These proteins co-localize at discrete ER foci, where LDs form in response to fatty acid loading^14,15^. Seipin is a ∼46-kDa protein with two transmembrane segments and a conserved ER lumenal domain that adopts an α/β-sandwich fold, a domain common in lipid-binding proteins^16–20^. Seipin assembles into a large oligomeric ring of 10–12 protomers (depending on species), with lumenal domains forming the base and transmembrane segments forming the walls of the cage-like complex^21,22^. LDAF1 is a ∼17-kDa protein with transmembrane segments of unknown structure^1,2^. Together, seipin and LDAF1 form a stoichiometric oligomeric complex of ∼700–850-kDa that defines the site of LD formation in cells^2^.

Although the LDAC is essential for normal LD formation, its molecular function has remained a mystery. A key limitation has been that prior studies of the LDAC largely relied on cellular systems, where LD formation is intertwined with metabolism, feedback regulation, and membrane dynamics, complicating mechanistic dissection. As a result, diverse molecular functions have been proposed for seipin alone and LDACs, including acting as a scaffold for TG synthesis enzymes^23^, concentrating lipids such as phosphatidic acid needed for TG synthesis^24–26^, modulating ER Ca^2+^ channels^27–29^, or facilitating TG accumulation at sites of LD formation^30–32^. However, direct evidence for these functions has been difficult to obtain *in vivo*, underscoring the need for reconstituted systems to define LDAC activity in isolation.

## The LDAC is necessary and sufficient for lipid droplet formation *in vitro*

We sought to determine if LDACs facilitate LD biogenesis by associating with TG molecules within the ER membrane. To test this hypothesis, we purified LDACs composed of seipin and LDAF1, as well as seipin alone, and an unrelated multitopic ER membrane protein (MBOAT7) as a control. Consistent with prior studies showing that LDAF1 stability depends on seipin in cells^2^, we were unable to recombinantly produce LDAF1 without seipin. We reconstituted the purified proteins into membranes containing 2.5 mol% TG—a concentration below the phase separation threshold—and included radiolabeled TG to trace lipid association. We then re-extracted the proteins with the mild detergent GDN and measured TG binding (Fig. 1a–c, Extended Data Fig. 1a–d). Only the LDAC, containing both seipin and LDAF1, but not seipin alone or MBOAT7, exhibited increased TG association. Increased association of LDACs, but not seipin alone, with TG also occurred at 6 mol% TG, indicating that the enhanced binding was independent of TG concentration in this range (Fig. 1d,e).

**Fig. 1.**
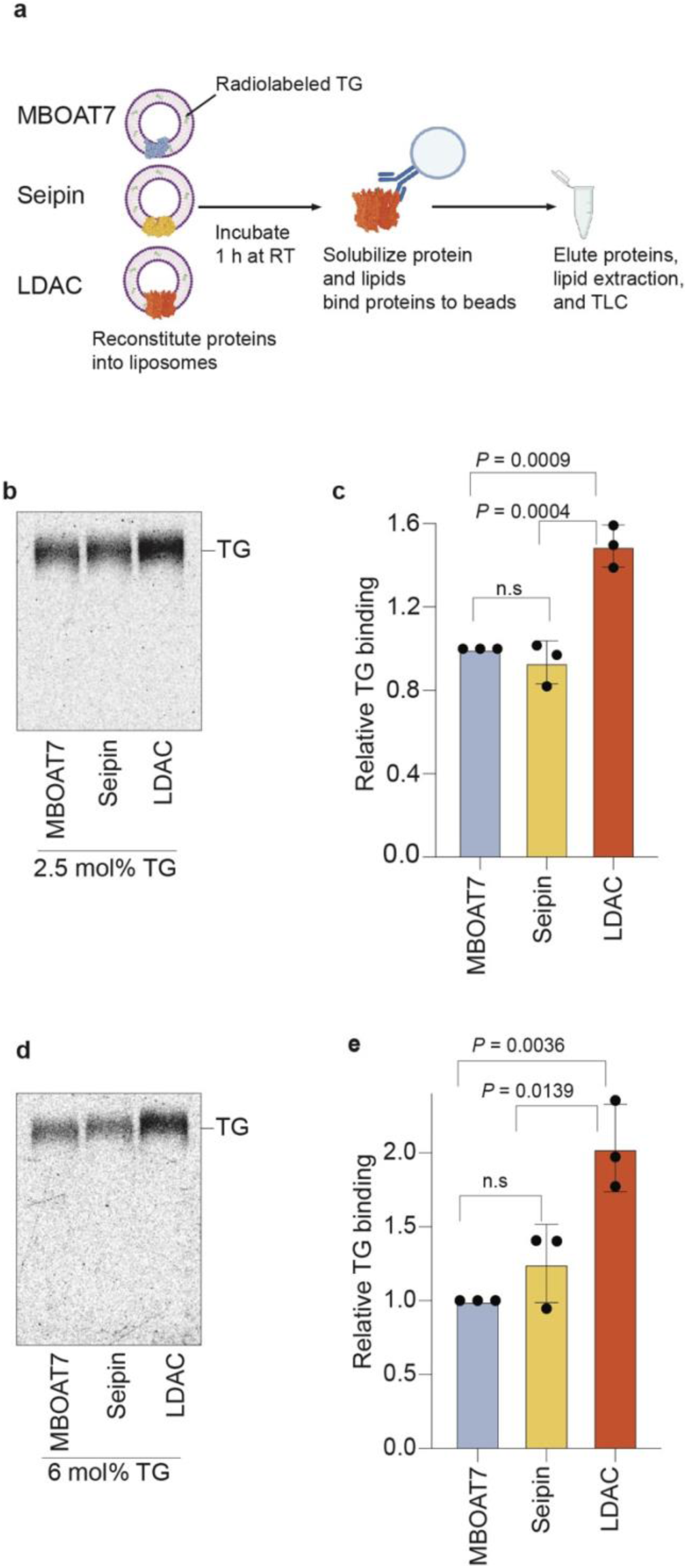
Liposome reconstitution reveals that the LDAC is necessary for TG association *in vitro*. **a,** Schematic representation of TG binding assay. Phospholipids depicted in purple, radiolabeled TGs in green, MBOAT7 in light blue, seipin in yellow, and LDAC in orange. **b,** Purified LDAC, but not seipin alone or MBOAT7 binds TG at 2.5 mol% TG. Elution of re-purified proteins was loaded on TLC to separate radiolabeled TG. **c,** Quantitation of signal intensity corresponding to TG in the TLCs. One-way ANOVA with Tukey’s post hoc test was performed (mean ± s.d., n=3). **d,** LDAC but no seipin alone binds TG at 6 mol% TG. TG binding assay with purified proteins at 6 mol% TG. Elution of re-purified proteins was loaded on TLC to separate radiolabeled TG. **e,** Quantitation of signal intensity correspond to TG in the TLCs. One-way ANOVA with Tukey’s post hoc test was performed (mean ± s.d., n=3).

To define the molecular function of the LDAC, we developed a minimal reconstitution system to study LD formation *in vitro*, independent of cellular regulation. Using giant unilamellar vesicles (GUVs) containing defined TG concentrations, we found that their bilayers composed of phosphatidylcholine (PC) accommodated TG up to 4 mol%. Above this concentration, TG molecules phase-separated, forming oil lenses within the membrane (Fig. 2a,b).

**Fig. 2.**
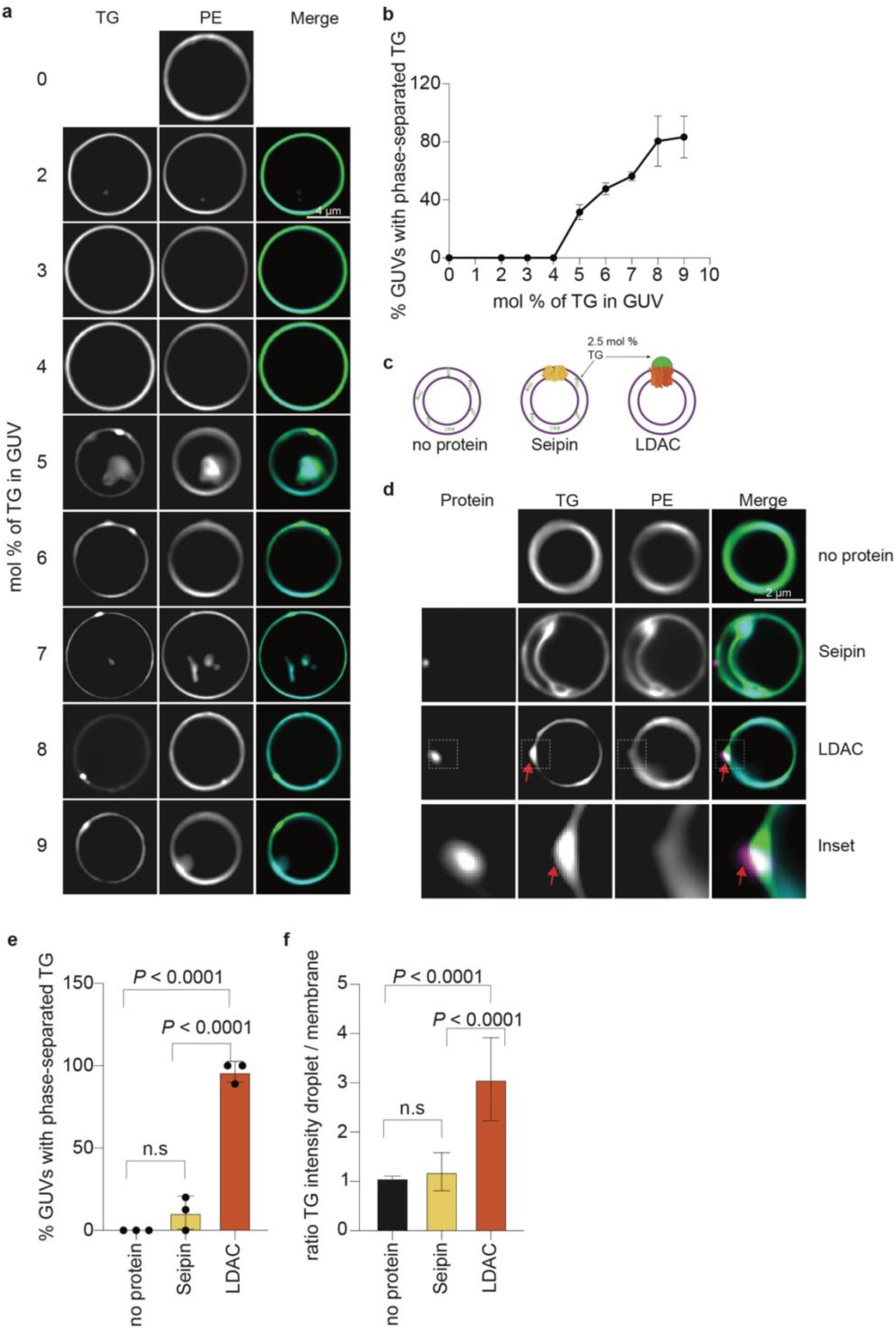
Biochemical reconstitution reveals that the LDAC is necessary and sufficient for LD formation *in vitro*. **a,** GUVs can accommodate up to 4 mol% TG. GUVs with varying TG concentration (mean ± s.d., n > 10 GUVs per condition, n=3) to assess spontaneous phase separation. **b,** Quantitation of phase-separated TG in a, (mean ± s.d., n=3). **c,** Schematic representation of GUV assay, soluble concentrations of TG were added to GUVs with no protein, seipin alone or LDAC. Phospholipids are labeled with PE-ATTO390, TG with top fluor TG, and protein with rhodamine. **d,** LDAC but not seipin alone triggers TG demixing in GUVs. Representative confocal images of GUVs showing TG (green), phospholipids (cyan), and protein signal (magenta). **e,** Quantitation of phase-separated TG in the different conditions. One-way ANOVA with Tukey’s post hoc test was performed (mean ± s.d., n=3). **f,** Quantitation of signal intensity from GUV images. A ratio between the signal in the phase-separated TG and the membrane was performed. One-way ANOVA with Tukey’s post hoc test was performed (mean ± s.d., n=3).

Next, we tested the hypothesis that LDAC association with TG promotes their separation into an oil droplet below the critical concentration of spontaneous phase separation in the membrane. We reconstituted fluorescently labeled LDACs or seipin alone into GUVs containing 2.5 mol% TG, along with trace amounts of fluorescent phosphatidylethanolamine (PE) to mark the bilayer and fluorescent TG to monitor neutral lipid behavior (Fig. 2c). As expected, TG remained uniformly distributed within the membrane in the absence of protein (Fig. 2d). Reconstitution of seipin alone, although previously proposed to promote TG demixing, had only a minimal effect under these conditions, with TG phase separation occurring in two of the 20 GUVs analyzed. In contrast, reconstitution of the LDAC containing seipin and LDAF1 into GUVs triggered TG demixing into a single, focused region, accompanied by depletion of TG from the surrounding membrane in 21 of the 22 GUVs analyzed (Fig. 2d–f, Extended Data Fig. 1e). The PE signal marking phospholipids remained evenly distributed, showing that the membrane remained unilamellar at sites of TG focus formation. Notably, the TG focus always colocalized with fluorescently labeled seipin, marking the position of the LDAC. This data demonstrates that the LDAC is the minimal complex necessary and sufficient to promote oil droplet formation *in vitro*.

## The LDAC forms a toroid-shaped assembly

To understand how the LDAC promotes oil-phase formation, we determined its structure by cryogenic electron microscopy (cryo-EM). We reconstituted LDAC into nanodiscs, vitrified, and imaged them by cryo-EM. From the resulting images, we generated an *ab initio* 3D reconstruction using high-quality 2D class averages and a set of decoy 3D models. Iterative heterogeneous refinement of these models resolved the architecture of the complex (Extended Data Fig. 2a,b).

The resulting cryo-EM map revealed the highest resolution in the lumenal domain of seipin (Extended Data Fig. 3a-e), with lower resolution but discernible density in the transmembrane regions of both seipin and LDAF1. To build a molecular model of the LDAC, we combined cryo-EM-guided model building with AlphaFold3 structural predictions^33^, using a minimal, functional fragment of LDAF1 (residues 45–135)^2^ in complex with seipin (residues 1–263) (Extended Data Fig. 2c). The fitted model resembled the reported architecture of the seipin transmembrane segments in the “A” conformation for yeast seipin^22^ (Fig. 3a). Together, seipin and LDAF1 formed a double-ring toroid structure with distinct architecture: an inner ring of tightly packed LDAF1 helices, and an outer ring of more loosely spaced seipin transmembrane segments that showed gaps within the plane of the membrane between adjacent seipin protomers (Fig. 3a). The lumenal domain matched the canonical α/β-sandwich structure but, owing to the presence of LDAF1, was rotated at the switch region connecting it to the transmembrane helices, shifting the domain toward the ER lumen (Fig. 3b, orange region). The two transmembrane helices of LDAF1 were resolved, contacting the hydrophobic helix of the lumenal domain of seipin and inserting into the center of the ring (Fig. 3c). On the lumenal side, the transmembrane helices of both seipin and LDAF1 proteins were capped by the α/β-sandwich domains of seipin. Evolutionary couplings^34^ between residues in seipin and LDAF1 (Extended Data Fig. 4a,b) were consistent with the structure of LDAC and found at positions where both proteins physically interact with each other (Fig. 3d and Extended Data Fig. 4a,b).

**Fig. 3.**
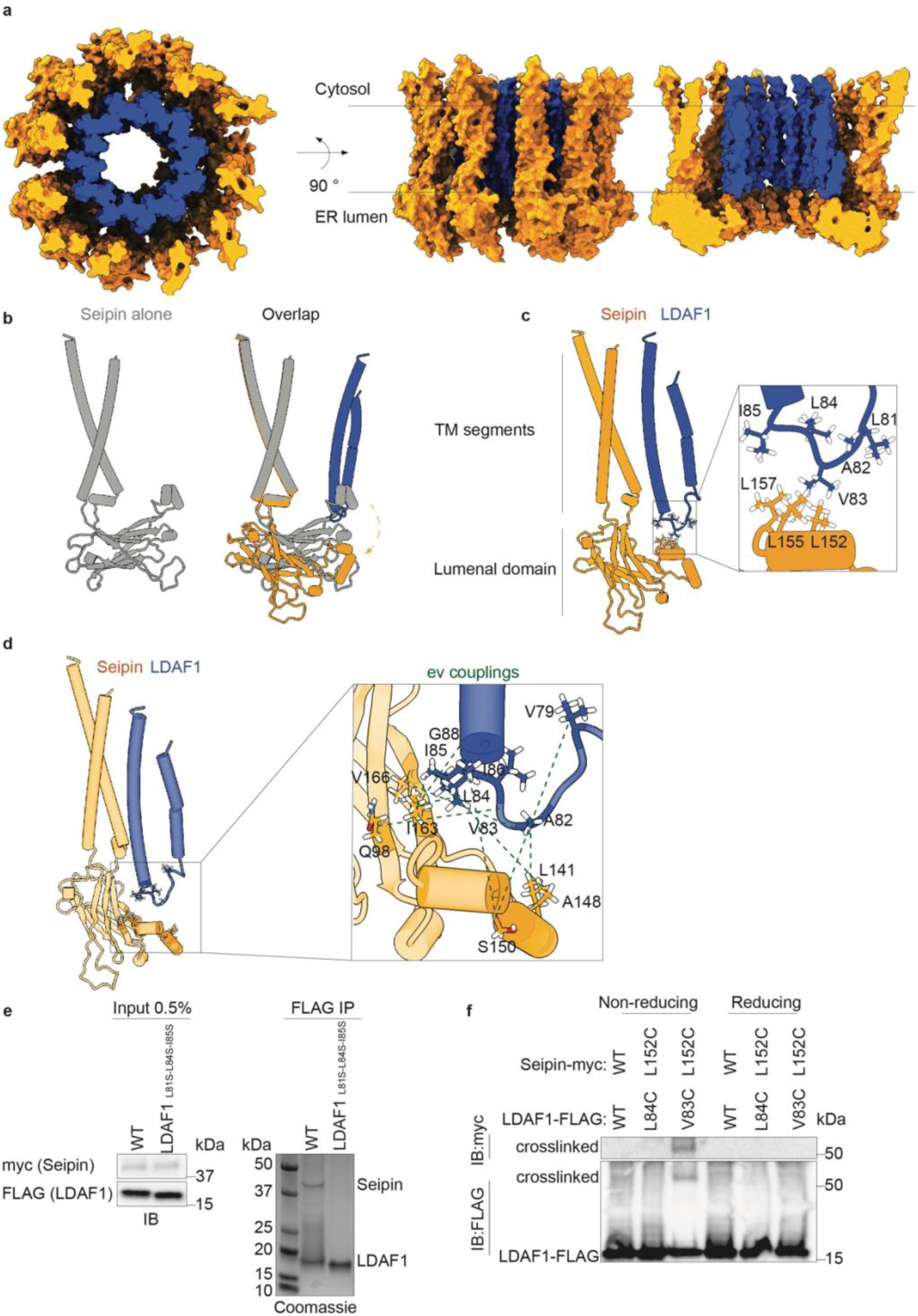
Structural analysis shows that the LDAC forms a toroid-shaped oligomeric assembly. **a,** Seipin and LDAF1 formed a double-ring toroid structure. Space filling model of the LDAC from a top and side views with the plane of the membrane **b,** Overlay of the structures of seipin alone or with LDAC. The lumenal domain of seipin undergoes a rigid-body movement (arrow). **c,** Cylinder representation of seipin and LDAF1. Magnified box shows details of the interacting residues between seipin and LDAF1. **d,** Evolutionary couplings between seipin-LDAF1 mapped onto the structure model. **e,** Mutations in LDAF1 at the interaction interface with seipin disrupted the binding. Immunoprecipitation of WT and mutant LDAF1. Immunoblot of the total lysate on the left, and Coomassie-stained SDS-page gel of resultant immunoprecipitation. **f,** Introduction of cysteine pair between LDAF1 Val83 and seipin Leu152 led to a crosslinked complex. Immunoprecipitation of WT and mutant complexes in non-reducing and reducing environment. Immunoblot of the precipitated complexes.

The interaction between seipin and LDAF1 appears to be mediated by hydrophobic contacts between seipin residues Leu152, Leu155, and Leu157 and LDAF1 residues Leu81, Val83, Leu84, and Ile85 at the interface of their respective helices (Fig. 3c). To test the functional relevance of these contacts, we mutated LDAF1 residues Leu81, Leu84, and Ile85 to serine. These substitutions disrupted complex formation, abolishing the interaction between LDAF1 and seipin (Fig. 3e). To test whether residues that are adjacent within our model are indeed physically associating with each other, we mutated LDAF1 residues Val83 and Leu84 and seipin residue Leu152 to cysteines and performed a cysteine crosslinking experiment. Incorporation of cysteines in LDAF1 Val83 and seipin Leu152 led to a crosslinked complex running at the expected size of 54 kDa. The crosslinking was reversed in a reducing environment (Fig. 3f).

## The LDAC facilitates lipid droplet formation by selectively concentrating TG

The LDAC structure prompted us to ask how the complex promotes neutral lipid phase separation within the membrane. Previous molecular dynamics (MD) simulations performed with seipin alone suggested that serine residues in a short hydrophobic helix of the lumenal domain projecting into the membrane may bind TG molecules^31,32^. In the LDAC structure, however, these residues do not insert into the membrane but instead are displaced outward from the membrane and face the interior of the seipin lumenal ring, raising questions about their function. Moreover, seipin alone did not bind TG or promote phase separation *in vitro* (Fig. 1a-e; Fig. 2 d,e; Extended Data Fig. 1a,e). This highlights the role of LDAF1 in facilitating TG phase separation.

To gain mechanistic insight into LDAC-mediated TG phase separation, we used the new structural data to perform MD simulations. We first simulated a single LDAF1 protomer and a TG molecule embedded in a membrane bilayer. In coarse-grain simulation (CG), the glycerol moiety of TG reproducibly bound and unbound LDAF1 at conserved serines Ser61 and Ser109 (Fig. 4a,b). To test how LDAC catalyzes TG lens formation, we simulated its behavior in a membrane containing PC and TG (Fig. 4c). We observed that the TG diffusion coefficient was 20-fold lower within the LDAC than in the bulk membrane (Fig. 4d). In contrast, seipin alone showed a 10-fold reduction in TG diffusion coefficient relative to that in the bulk membrane.

**Fig. 4.**
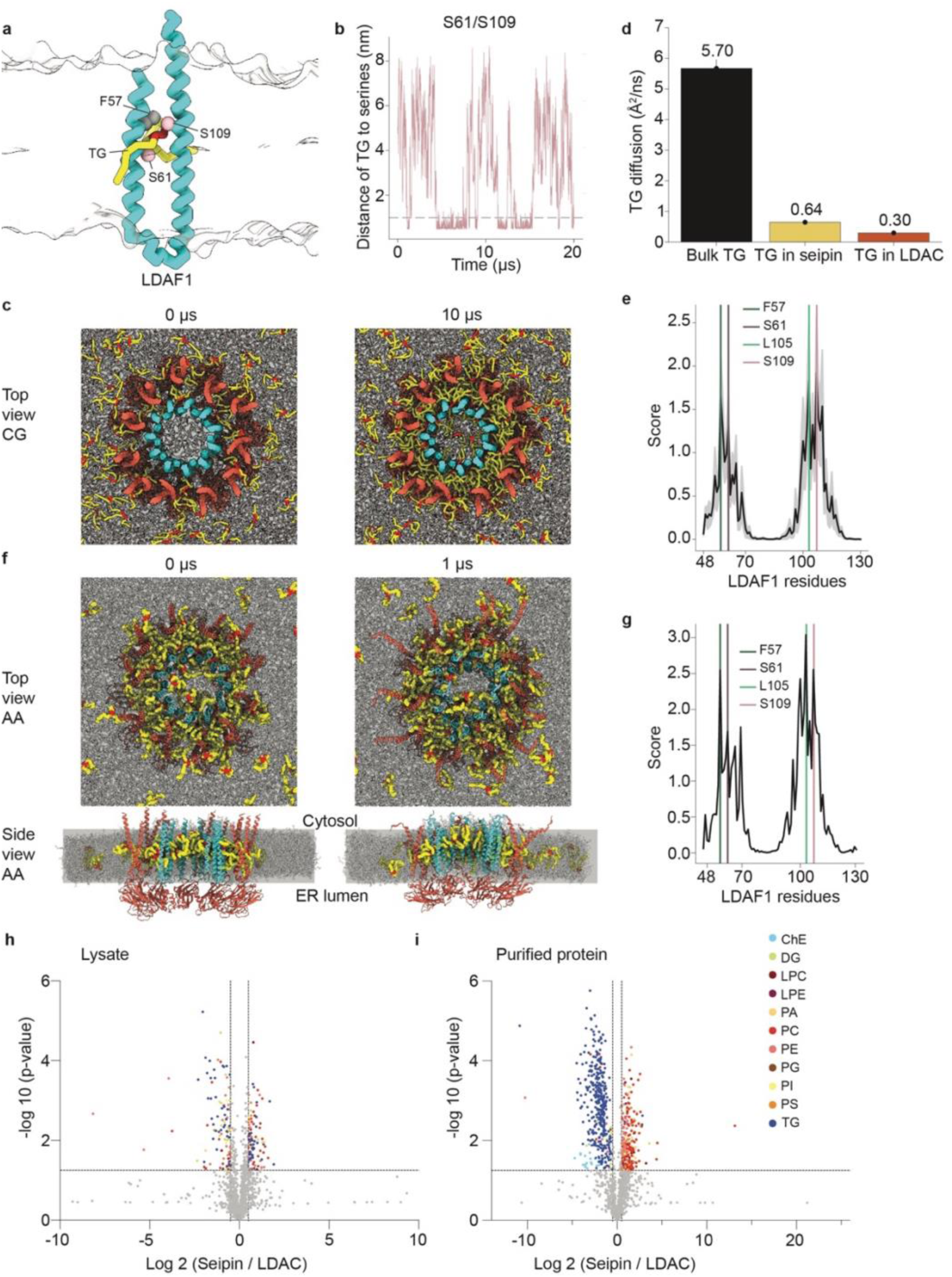
MD simulations show TG binding to LDAF1 and accumulation in the central toroid-shaped cavity formed by seipin and LDAF1. **a,** A representative CG frame illustrating TG binding to LDAF1. **b,** Serines 61 and 109 in LDAF1 bind TG. Distances between the central glycerol group of TG and key serine residues. **c,** TG accumulates in the central toroid-shaped cavity of the LDAC in CG simulations. Top view of the CG simulations at 0 and 10 microseconds with 5 mol% TG. Seipin is shown in red, LDAF1 in cyan, TG tails are shown in yellow, and TG glycerol groups in red. **d,** TG diffuses slower in the LDAC complex than seipin alone or bulk membrane. Diffusion coefficients of TG in the bulk membrane, within seipin, and the LDAC. **e,** Key residues in LDAF1 interact with TG in CG simulations. Interaction score analysis between LDAF1 and TG. Vertical lines mark four conserved residues that were mutated in the experimental mutagenesis study. **f,** TG accumulates in the central toroid-shaped cavity of the LDAC in AA simulations. Top and side view of the AA simulation at 0 and 1 microseconds with 5 mol% TG. Seipin is shown in red, LDAF1 in cyan, TG tails are shown in yellow, and TG glycerol groups in red. **g,** Key residues in LDAF1 interact with TG in AA simulations. Interaction score analysis between LDAF1 and TG calculated from the AA trajectory. **h,** Expression of LDAC or seipin did not alter the lipidome. Volcano plot of total cell lysate lipidome illustrating different lipid species; p-value was calculated in two sample t-test. **i,** LDAC is enriched in neutral lipids and reduced association with phospholipids. Volcano plot of isolated protein lipidome illustrating different lipid species, p-value was calculated in two sample t-test.

As the CG simulations progressed, TG molecules, initially dispersed uniformly in the bilayer, coalesced between the inner LDAF1 and outer seipin transmembrane rings, forming a stable oil phase within the toroid (Fig. 4c). Within this space, TGs preferentially interacted with conserved LDAF1 residues (i.e., Phe57, Ser61, Leu105, and Ser109) (Fig. 4e). In agreement with these interactions, alanine substitution of these residues reduced TG binding to the complex *in vitro* (Extended Data Fig. 5a-d). These findings suggest that the interactions between TG and the conserved residues are critical, and the LDAC promotes phase separation more effectively than seipin alone, due to stronger and more localized TG binding.

To understand atomistic protein-lipid interactions, we backmapped a phase-separated CG structure to all-atom (AA) resolution and conducted additional simulations^35^. The oil lens was maintained in a double-ring toroid structure (Fig. 4f), and the key interactions between LDAF1 residues and TG (Fig. 4g) were consistent overall with the CG data (Fig. 4e). The diffusion coefficients of TG in the membrane and within the LDAC were calculated to be 1.1 A²/ns and 0.2 A²/ns, respectively, capturing a 5.5-fold reduction in TG diffusion within the LDAC. These simulations without any restraints on protein movement in the LDAC suggested a protein conformational change. The N-terminal transmembrane segments of seipin, opened up, and the lumenal domain shifted toward the ER membrane, mimicking the initial stage of membrane deformation and positive curvature formation, as shown in previous CG simulations^36^ and cryo-electron tomography^37^ (Fig. 4f, side view). In contrast, this motion was not observed when TG was not phase-separated in AA simulations of the LDAC (Extended Data Fig. 6a,b).

These findings suggest that the LDAC catalyzes LD formation by binding TG while limiting phospholipids within the LDAC core, thereby promoting TG–TG interactions and phase separation. This model predicts that LDACs in cells should preferentially associates with TGs over phospholipids. To test this, we purified seipin alone or LDACs from cells and analyzed their associated lipids by mass spectrometry–based lipidomics. In agreement with the model, LDACs selectively enriched TGs and showed less association with phospholipids than seipin alone (Fig. 4h,i; Extended Data Fig. 7,8). Importantly, expression of either LDAC or seipin did not significantly alter the bulk cellular lipidome, indicating that lipid enrichment reflected selective association rather than changes in lipid synthesis or metabolism.

## Discussion

Collectively, our findings provide a model for the molecular mechanism of LD formation within the ER bilayer. At TG concentrations below the threshold for spontaneous phase separation (e.g., ≤4 mol% *in vitro*), LDACs are necessary and sufficient to initiate TG demixing within a phospholipid membrane bilayer. The multimeric, membrane-embedded LDAC forms a central chamber, enclosed by inner and outer transmembrane rings. This architecture allows TG entry and interaction with LDAF1 residues, slowing the diffusion of TG and promoting TG phase separation and LD formation. Given that seipin is implicated in the formation of LDs composed pf other neutral lipids, such as sterol esters^38,39^, this mechanism likely reflects a general principle of LDAC-mediated neutral lipid phase separation.

While these findings define a core mechanism for LDAC-mediated LD biogenesis, several questions remain. In cells, seipin and LDAF1 may have additional functions beyond their minimal biochemical activity. The more severe phenotypes in seipin than LDAF1-deficient cells^2,40^ further suggest that seipin also acts independently (e.g., to maintain ER–LD connections during LD growth^18,41^, a process likely essential for lipid storage and turnover in cells). In yeast, the seipin orthologue Sei1 may form a similarly functioning LDAC with Ldb16 and Ldo45^14,15^.

In specialized cell types, such as adipocytes, the core LDAC activity may be further modulated by accessory proteins, including adipogenin^42^ and other lipid metabolic factors. These observations point to additional layers of regulation superimposed on the core mechanism described herein.

## Methods

### Protein expression and purification

The LDAC and seipin were expressed in HEK293 Gnti^−^ suspension cells. Cells were cultured in Free Style medium (Thermo Fisher Scientific, #12338026) at 37°C under 8% CO_2_ and 80% humidity in a Multitron-Pro shaker at 125 rpm. When cell density reached 2 x 10^6^ cells per mL, pCAG-LNK plasmids were transfected. 1 mg of plasmid was pre-mixed with 3 mg of polyethamine (Polysciences, #23966-1) in Opti-MEM medium for 30 min at room temperature. At 16 h after transfection, cells were supplemented with 10 mM sodium butyrate to boost protein expression. Cells were collected 48 h after transfection, snap frozen in liquid nitrogen and stored at -80°C.

Protein purification was performed at 4°C. Cell pellets were resuspended in 400 mM NaCl, 50 mM Tris, pH 8.0, 0.5 mM EDTA supplemented with complete protease inhibitor tablet, EDTA free (Roche, #5056489001). Cells were lysed with a dounce homogenizer, and the lysate was supplemented with 1% detergent. After 2 h rotating at 4°C, the cell lysate was spun at 50.000 x g for 45 min. The supernatant was incubated with anti-FLAG M2 resin (Sigma-Aldrich, #A2220) for 2 h at 4°C. Resin was collected and washed with 10 column volumes with buffer A (400 mM NaCl, 50 mM Tris, pH 8.0, 5 mM MgCl_2_) supplemented with 0.1% detergent. Proteins were eluted with buffer A supplemented with 0.05% detergent and 0.2 mg/mL of 3xFLAG peptide (Sigma-Aldrich, #F4799). The elution fraction was concentrated and further purified by size-exclusion chromatography on a Superose 6 3.2/300 increase column equilibrated with 150 mM NaCl, 50 mM HEPES, pH 7.4, 5 mM MgCl_2_, 0.05% detergent. Peak fractions were collected and concentrated. For cryo-EM analysis, the LDAC was reconstituted into nanodiscs. Purified LDAC in detergent was mixed with MSP2N2 and POPG (Avanti, #840457) at a 1:2:130 molar ratio. After incubation for 30 min at room temperature, 20 mg of Bio-Beads (Biorad, #152-8920) were added, and the mixture was incubated at 4°C with gentle agitation for 1 h, followed by addition of 20 mg Bio-Beads and another hour incubation at 4°C. Finally, a last batch of 20 mg of Bio-Beads was added and incubated at 4°C overnight. The sample was then incubated with MS(PEG)12 methyl-PEG-NHS-ester (Thermo Fisher Scientific, #22685) at 1:10 molar ratio for 2 h at 4°C to reduce aggregation of particles on the grid. The reconstitution solution was filtered through a 0.22-μm filter (Corning, #8160) and further purified by size-exclusion chromatography.

### Protein reconstitution into liposomes and *in vitro* TG binding assay

A lipid mixture of 97.5 mol % DOPC and 2.5 mol % [^14^C] triolein was prepared in chloroform (Avanti Polar Lipids and American Radiolabeled Chemicals). Lipids were dried under a N_2_ stream and then placed under vacuum in a desiccator for 2 h. The lipid film was resuspended in buffer (150 mM NaCl, 20 mM HEPES, pH 7.4), and placed in a sonicator bath for 10 min at RT. The mixture was then extruded through a polycarbonate membrane of 200-nm pore size (Whatman, #10417006). Extruded lipids were incubated with 450 μM detergent for 1 h at RT with head-to-head rotation.

Lipids and proteins were mixed in a 1:50 molar ratio and incubated at RT for 1 h with head-to-head rotation. Then, Bio-Beads were added in three batches of 20 mg each with a 1-h incubation time for the first two batches and overnight incubation for the last one at 4°C. Successful reconstitution was assessed by analyzing flotation on a density gradient. Proteins incorporated into the liposomes floated to the top of the gradient.

Proteins reconstituted in liposomes were left to equilibrate for 1 h at RT with head-to-head rotation, then 0.5% detergent was added and incubated at 4°C for 1.5 h, followed by incubation with anti-FLAG M2 resin for 2 h at 4°C. The flow-through was collected, and protein was eluted with 0.2 mg/mL of 3xFLAG peptide. Lipids from flow-through and proteins were extracted by adding 6 mL of chloroform:methanol 2:1 (v:v) and incubated overnight at 4°C with rotation. Then, 1.5 mL of ultra-pure water was added, and spun for 20 min at 1,000 rpm. Bottom phase was collected with a glass pipette, and the lipids were dried under a N_2_ stream. Lipids were separated on a thin layer chromatography (TLC) with hexane:diethyl ether:acetic acid (80:20:1) solvent system. TLC plates were exposed to phosphor imaging screen overnight and develop by Typhoon phospho-imager.

### Protein labeling with maleimide

Proteins were purified as described above. After elution, proteins were reduced with 1 mM TCEP. Rhodamine Red C2 Maleimide (Thermo Fisher Scientific, #R6029) was added, and labeling was performed, following the manufacturer’s instructions. Then, proteins were further purified by size-exclusion chromatography on a Superose 6 3.2/300 increase column equilibrated with 150 mM NaCl, 50 mM HEPES, pH 7.4, 5 mM MgCl_2_, and 0.05% detergent.

### Protein incorporation into GUVs

A lipid mixture of 96.5 mol % DOPC, 1.0 mol % ATTO390PE (ATTO-TEC, #390-161), 2.0 mol % triolein and 0.5 mol % TopFluorTriolein (Avanti Polar Lipids, #810298C) was prepared in chloroform. Also, 19 μM detergent was added to the mixture to aid with protein incorporation as described^43^. Lipids in chloroform with detergent were spread on indium tin oxide–coated glass slides. GUVs were prepared by electroformation^44^ in 600 mM sucrose with a Vesicle Prep Pro (Nanion) with the following settings: frequency 10 Hz, amplitude 1.4 V, temperature 23°C for 1 h. GUVs were collected and incubated with protein in detergent. The mixture was incubated for 30 min at RT with head-to-head rotation. Then, Bio-Beads were added in three batches of 20 mg each with 1 h incubation time for the first two batches and overnight incubation for the last one at 4°C.

### Fluorescence microscopy

Spinning-disk confocal microscopy was conducted using a Nikon Eclipse Ti2 inverted microscope with Perfect Focus, CSU-X1 spinning disk confocal head (Yokogawa), ORCA-fusion BT scientific complementary metal-oxide semiconductor (sCMOS) camera (Hamamatsu), and NIS-Elements software (Nikon). Images were acquired through a 60x Apo TIRF 1.49 NA objective with SoRa function. Image pixel size was 0.03 μm/px. Blue, green and red fluorescence was excited by 405-, 488-, and 561-nm lasers. Multicolor images were acquired sequentially.

### Image analysis

ImageJ^45^ was used to adjust the contrast and convert to 8-bit microscopy images. Phase separation of TG in GUV assays was quantified manually with ImageJ. Fluorescence intensity on LD and GUV bilayer was quantified to calculate enrichment of fluorescence on LDs.

### Cryo-electron microscopy sample preparation and data acquisition

Purified LDAC (2–3 μL) in nanodisc was applied to Quantifoil holey carbon grids (Au R1.2/1.3; 400 mesh) that were glow-discharged for 30 s. Proteins were concentrated to 1–2 mg/mL. Grids were blotted for 6 s with a Whatman #1 paper at 4°C with ∼ 95% humidity and plunge frozen in liquid ethane cooled with liquid nitrogen using a Vitrobot Mark IV system (Thermo Fisher Scientific). Cryo-EM data was collected on a Titan Krios electron microscope (Thermo Fisher Scientific) at the New York Structural Biology Center, operated at 300 kV, with Gatan K3 imaging system collected with a physical pixel size of 1.083Å per pixel. Videos were collected using Leginon^46^ with an accumulated electron exposure of 51.29 e^-^ /Å^2^. A total of 13,063 images were collected at a nominal defocus range of 0.8–2.5 μm under focus.

### Cryo-electron microscopy image processing

Processing was conducted with cryoSPARC^47^. Drift and beam-induced motions were corrected with patch motion correction, and the contrast transfer function was estimated with patch CTF estimation. Particle picking was performed with Topaz^48^, extracted with a box size of 128 pixels, and then used for calculation of initial 2D class averages. The best classes were used for *ab initio* reconstruction (C1 symmetry), and then heterogenous refinement (C1 symmetry) using the Topaz particle stack as input and eight reference maps, seven decoy noise maps and one very good reference map. The decoy maps were generated by running *ab initio* reconstruction and killing the job after the first iteration was completed. This process was done one more time, resulting in one class where LDAF1 was visible with a resolution of 6.41Å. Classes were only the lumenal domain was resolved were selected and further refine using non-uniform refinement (C11 symmetry) resulting in a 3.2Å resolution map (FSC = 0.143 criterion), and in a local resolution range of 2.6–3.2Å as computed by local resolution estimation in cryoSPARC.

### Model building

EM maps and models were inspected in ChimeraX^49^. Model of the lumenal domain was built in COOT^50^ starting from the high-resolution region and iteratively refined in PHENIX^51^ real-space refinement procedure, followed by visual inspection in COOT. This iterative process was repeated until the model reached optimal geometrical statistics as evaluated by MolProbility^52^.

### Immunoprecipitation of purified proteins for lipidomics

Purification was performed as described above with the following differences. Pellets were resuspended in buffer A 150 mM NaCl, 50 mM HEPES, pH 7.4, and 0.5 mM MgCl_2_, supplemented with complete protease inhibitor tablet, EDTA free. Cells were lysed with a dounce homogenizer, and the lysate was supplemented with 1% detergent. The supernatant was incubated with anti-FLAG M2 resin for 2 h at 4°C. Resin was collected and washed with 10 column volumes with buffer A supplemented with 0.1% detergent. Proteins were eluted with buffer A supplemented with 0.05% detergent and 0.2 mg/mL of 3xFLAG peptide.

Lipids of the purified protein or the cell lysates were extracted by adding 6 mL of chloroform:methanol 2:1 (v:v) and incubated over night at 4°C with rotation. Then, 1.5 mL of ultra-pure water was added and spun for 20 min at 1,000 rpm. The bottom phase was collected with a glass pipette, and the lipids were dried under a N_2_ stream. The dried lipid film was reconstituted in 150 µL of 65:30:5 (isopropanol:acetonitrile:water) solution for lipidomic analysis.

### LC-MS/MS lipidomic analysis

Lipids were separated using ultra-high-performance liquid chromatography (UHPLC) coupled with tandem mass spectrometry (MS/MS). Briefly, UHPLC analysis was conducted on a C30 reverse-phase column (Thermo Acclaim C30, 2.1 x 150 mm, 2.6 μm) maintained at 50°C and connected to a Vanquish Horizon UHPLC system (S/N:6516208), along with an Orbitrap Exploris 240 MS (Thermo Fisher Scientific, S/N:MM10585C) equipped with a heated electrospray ionization probe (HESI). 5 μL of each sample was injected, with separate injections for positive and negative ionization modes. Mobile phase A included 40:60 water:acetonitrile with 10 mM ammonium formate and 0.1% formic acid, and mobile phase B consisted of 90:10 isopropanol:acetonitrile with the same additives. The chromatographic gradient involved: Initial isocratic elution at 30% B from -3 to 0 minutes, followed by a linear increase to 43% B (0–2 min), then 55% B (2–2.1 min), 65% B (2.1–12 min), 85% B (12–18 min), and 100% B (18–20 min). Holding at 100% B from 20–25 min, a linear decrease to 30% B by 25.1 min, and holding from 25.1–28 min. The flow rate was 0.26 ml/min. Mass spectrometer parameters were ion transfer tube temperature, 300°C; vaporizer temperature 275°C; Orbitrap resolution MS1, 120,000, MS2, 30,000; RF lens, 70%; maximum injection time 50 ms, MS2, 54 ms; AGC target MS1, standard, MS2, standard; positive ion voltage, 3250 V; negative ion voltage, 2500 V; Aux gas, 10 units; sheath gas, 40 units; sweep gas, 1 unit. HCD fragmentation, stepped 15%, 25%, 35%; data-dependent tandem mass spectrometry (ddMS2) cycle time, 1.5 s; isolation window, 1 m/z; microscans, 1 unit; intensity threshold, 1.0e4; dynamic exclusion time, 2.5 s; isotope exclusion was enabled. Full-scan mode with ddMS2 at m/z 250–1700 was performed. EASYIC TM was used for internal calibration. The raw data were search and aligned using LipidSearch 5.1 with the precursor tolerance at 5 ppm and product tolerance at 8 ppm. Further data post-processing filtering and normalization were performed using an in-house developed app, Lipidcruncher^53^. All semi-targeted quantifications were done using area under the curve was normalized to the area under the curve for respective internal standards and amount of BCA (#23225, Thermo Fisher Scientific) protein measurements.

### Cysteine crosslinking

Plasmids with the incorporated cysteines pairs were transfected. Cells were spun at 1000 x g for 5 min and resuspended in PBS supplemented with 500 µM 4,4′-dithiopyridine (Aldrithiol-4, Sigma-Aldrich, # 143057). After 30 min incubation at 37°C in a shaker, the reactions were quenched with 200 mM N-ethylmaleimide for 15 min on ice. Cells were then pelleted and flash frozen in liquid nitrogen. Immunoprecipitation was performed as described above.

### Immunoblotting

Protein concentration was determined, and cell lysates were mixed with Laemmli buffer (Bio-Rad, #1610747) and heated for 5 min at 65°C prior to SDS-PAGE. Gels were transferred to Immuno-Blot PVDF membranes (Bio-Rad, #1620177) with 1x Tris/glycine transfer buffer (Bio-Rad, #1610771) and methanol for 90 min at 90 V at 4°C. Membranes were blocked with 5% non-fat dry milk (Santa Cruz Biotechnology, #sc-2325) at room temperature for 60 min, and primary antibody was incubated overnight at 4°C with shaking. Membranes were washed for 15 min with Tris buffer saline with tween (TBS-T) and then incubated with appropriate HRP-conjugated secondary antibodies (Santa Cruz Biotechnology) for 60 min at RT prior to development with chemiluminescence Super Signal Pico or Dura reagent (Thermo Fisher Scientific).

### Plasmid construction

Restrictions enzymes were from New England Biolabs. Synthetic DNAs (gBlocks) and primers were acquired from Integrated DNA Technologies. Point mutations were done with QuickChange XL (Agilent Technologies, #200517). pCAG-LNK vector was modified from pCAGEN (addgene, #11160) as described^54^. Generation of the seipin-LDAF1 expression vectors for protein purification were generated as described^54^.

### Molecular dynamics simulations

MD simulations were conducted using GROMACS 2024^55,56^. AA simulations utilized the CHARMM36m protein force field^57^ and the CHARMM36 lipid force field^58^, and CG simulations employed the Martini2 force field^59–61^. The AA TG model, designed to replicate the experimental interfacial tension, was utilized^62^. Each chain in the CG TG model consists of one Na bead, four C1 beads, and one C3 bead^63^. During the CG simulations, backbone beads were constrained with a positional restraint constant of 1000 kJ/mol/nm² to avoid collapse between transmembrane segments, driven by overestimation of protein-protein interactions. The simulations were carried out at a temperature of 310 K and a pressure of 1 bar. For all-atom simulations, the Nose-Hoover thermostat^64,65^ and the Parrinello-Rahman barostat^66^ were applied, whereas the V-scale and C-scale methods^67,68^ were used for Martini simulations. Nonbonded interactions were truncated at 1.1 nm in CG simulations, and in AA simulations, they were force-switched between 1.0 and 1.2 nm. The initial configurations for the MD simulations were generated using the Multiscale Simulation Tool (mstool)^69,70^ and simulated as outlined in Table 2. The initial structure of the TG-containing AA simulation was derived by backmapping the CG structure, which had already been simulated for 10 µs and thus contained TG nucleation within the complex.

### Simulation analysis

A dissociation constant was calculated by

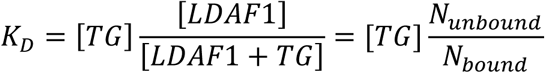

Here, N_unbound_ and N_bound_ denote the number of frames in which TG is unbound and bound to LDAF1, respectively.

Diffusion coefficient was calculated by

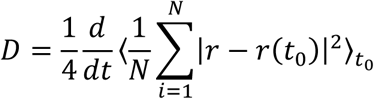

Where *N* is the number of molecules and *r* represents the 2D-XY coordinates of the central glycerol group of each TG molecule. The calculation was performed separately for two TG categories: molecules located far from the protein, and those positioned within seipin or the seipin– LDAF1 complex.

Interaction score was calculated by

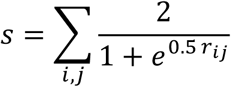

Where *r_ij_* is the distance between TG bead *i* and protein bead *j*. For each residue, the summation is performed over all TG beads and all protein beads belonging to that residue. The AA simulation was first mapped into the CG resolution before computing the score. Simulations were analyzed with MDAnalysis^71^.

### Statistical analysis

Three biological replicates were performed for all experiments. For the GUV assays ∼ 20 GUVs per condition were analyzed. To determine significance one-way analysis of variance (ANOVA) was performed followed by Tukey’s test. GraphPad Prism v.10 software was used.

---

**Extended Data Fig. 1.**
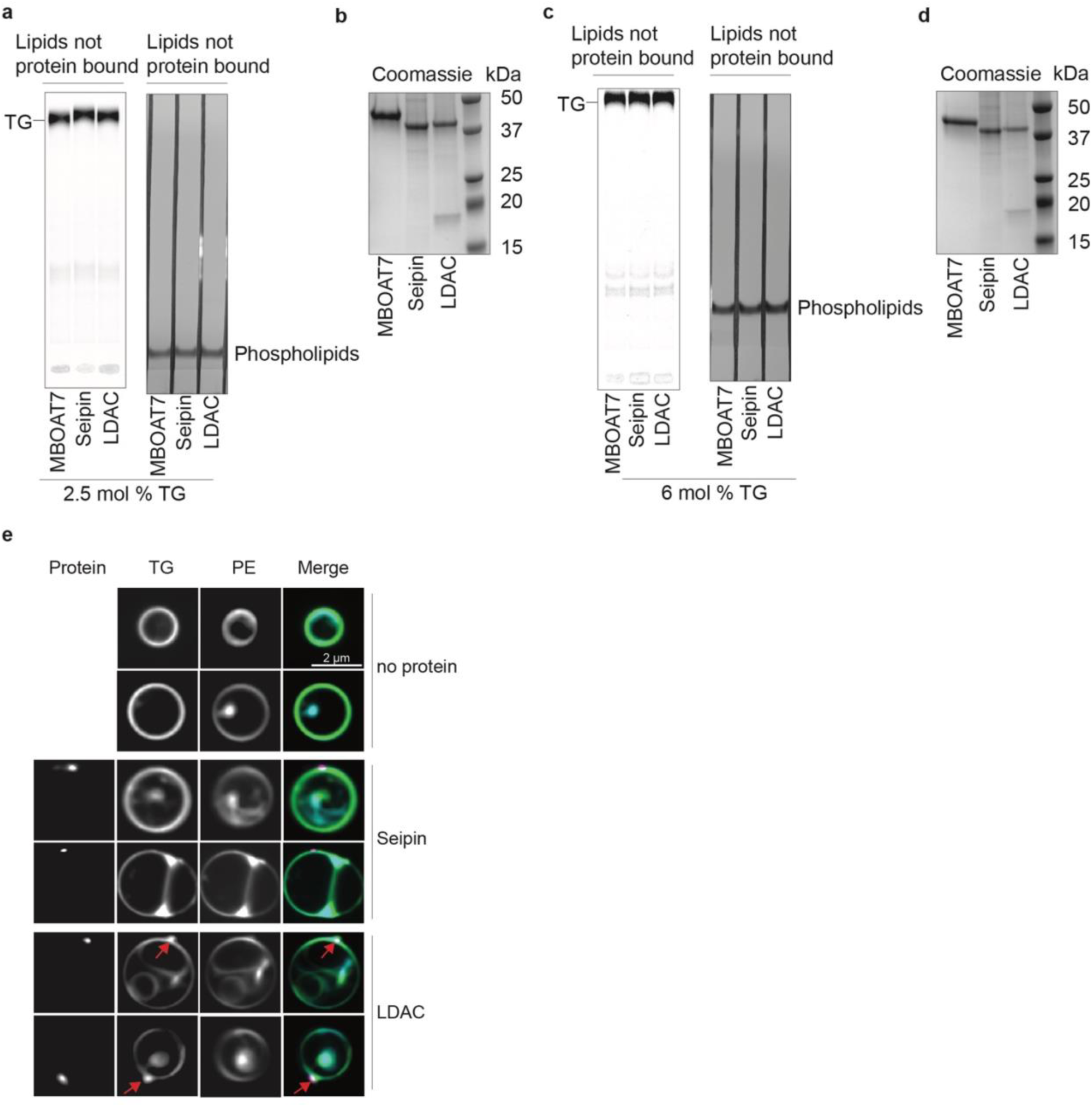
Biochemical reconstitution reveals that LDAC is necessary and sufficient for LD formation *in vitro*. **a,** All samples have equal amounts of phospholipids and TG as starting material. Flow-through of re-purified proteins from Fig 1b, analyzed by TLC. Radiolabeled TG was detected, as well as phospholipids that were stained with iodine vapor. **b,** Equal amount of proteins were used in each condition. Coomassie-stained SDS-page gel of purified proteins used in Fig 1b. **c,** All samples have equal amounts of phospholipids and TG as starting material. Flow-through of re-purified proteins from Figure 1 d, loaded on TLC. Radiolabeled TG was detected, as well as phospholipids that were stained with iodine vapor. **d,** Coomassie-stained SDS-page gel of purified proteins used in Figure 1 d. Equal amount of proteins were used in each condition. **e,** LDAC but no seipin alone triggers TG demixing in GUVs. Representative confocal images of GUVs showing TG (green), phospholipids (blue), and protein signal (magenta).

**Extended Data Fig. 2.**
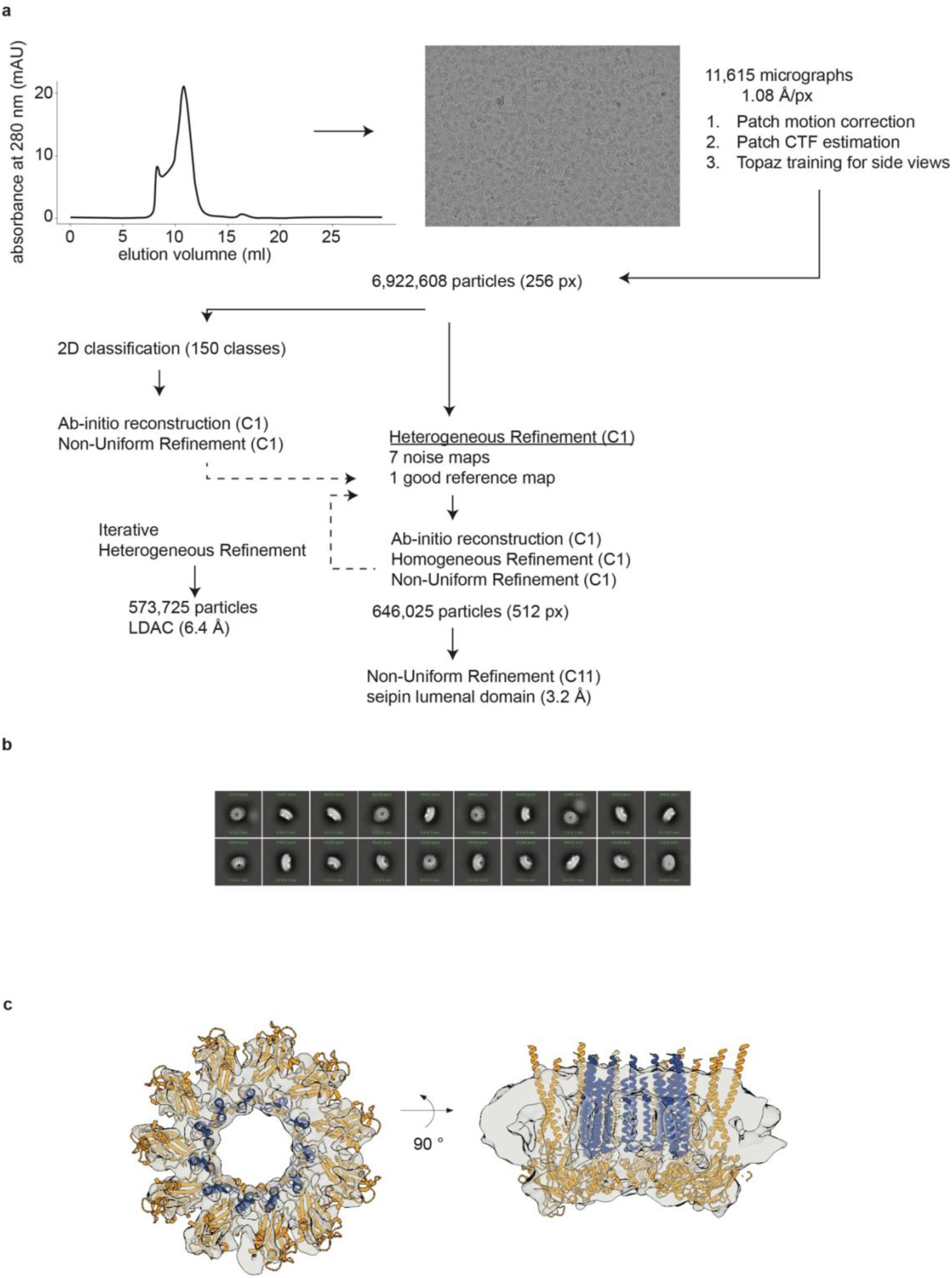
Cryo-EM processing workflow. **a,** Size-exclusion chromatography of LDAC in nanodisc shows a monodisperse peak. Representative micrograph and cryo-EM workflow. **b,** Representative 2D classes. **c,** Cryo-EM density map of the LDAC in nanodisc, with a fit of the model in the densities (seipin aa 1–263; LDAF1 aa 45–135).

**Extended Data Fig. 3.**
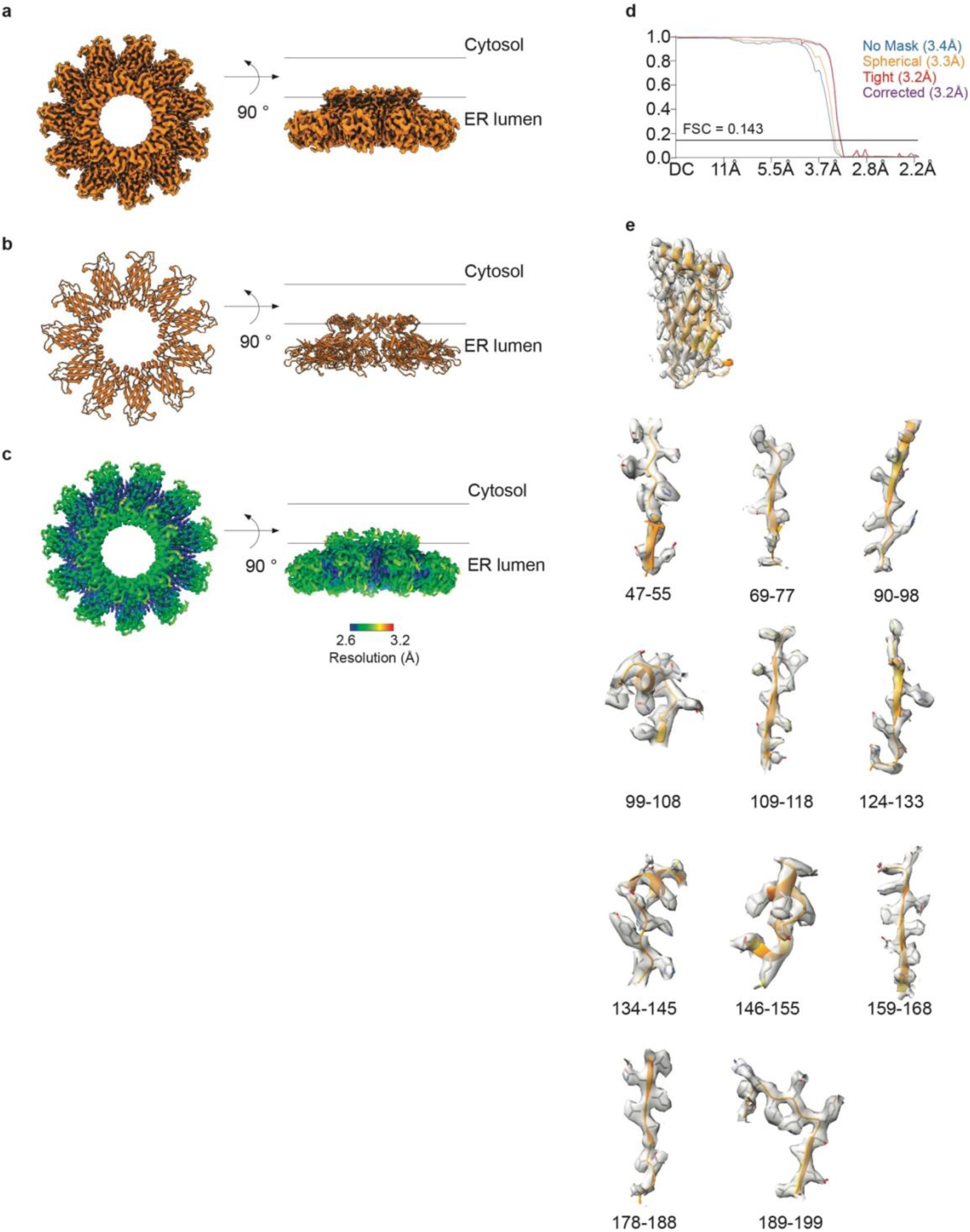
Single-particle cryo-EM analysis of seipin lumenal domain. **a,** EM map of seipin lumenal domain from a top view and on the membrane plane. **b,** Atomistic model of the seipin lumenal domain seen from the top and in the membrane plane. **c,** Local resolution mapped onto EM density map. **d,** FSC curves: gold standard FSC curve between two half maps with indicated resolution at FSC = 0.143. **e,** Superimposed cryo-EM densities from sharpened map with atomistic model.

**Extended Data Fig. 4.**
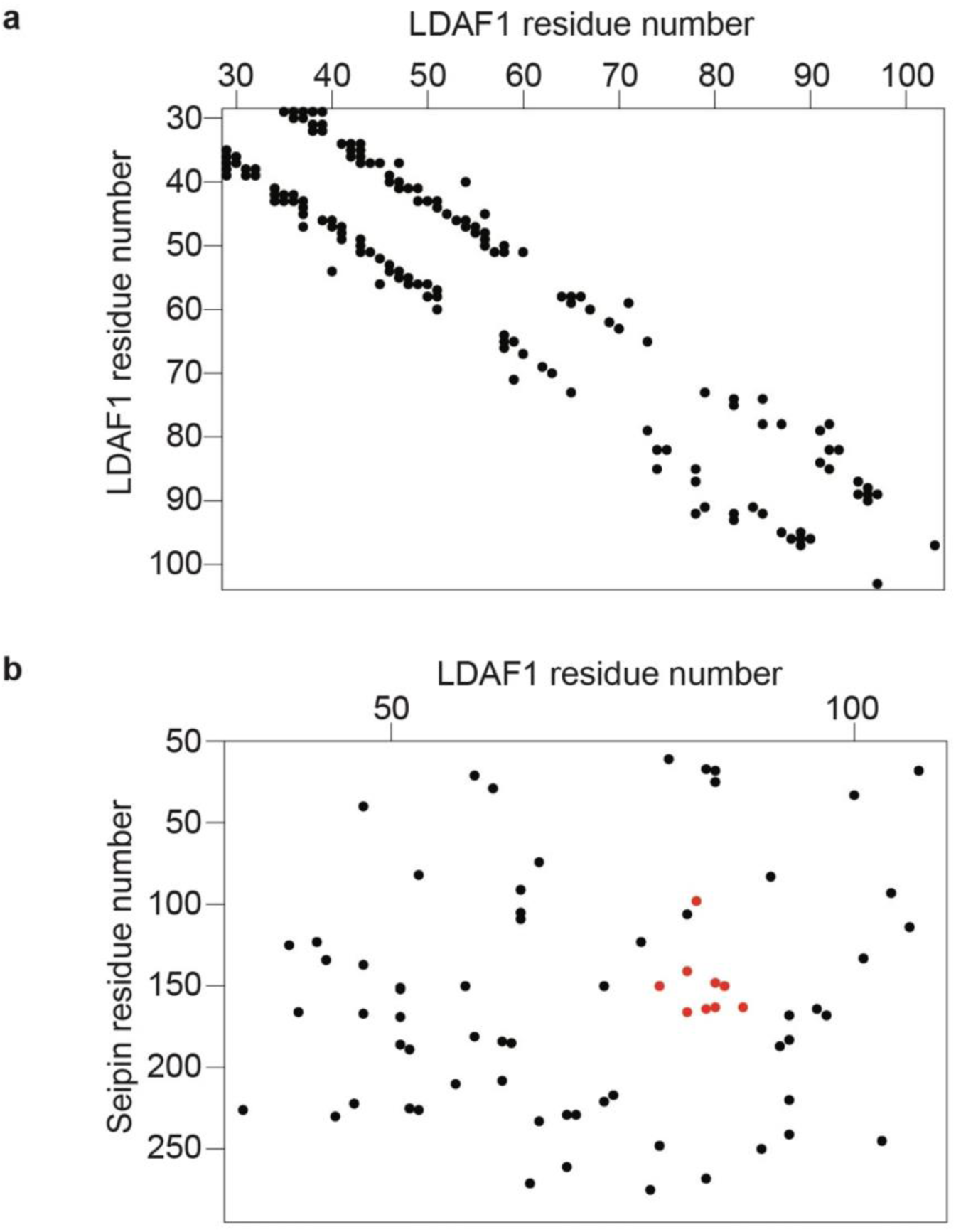
Evolutionary couplings of seipin and LDAF1 reveals co-evolving residues. **a,** Evolutionary couplings residues in LDAF1. **b,** Evolutionary couplings between seipin-LDAF1 in red couplings mapped into the structure in Fig. 3d.

**Extended Data Fig. 5.**
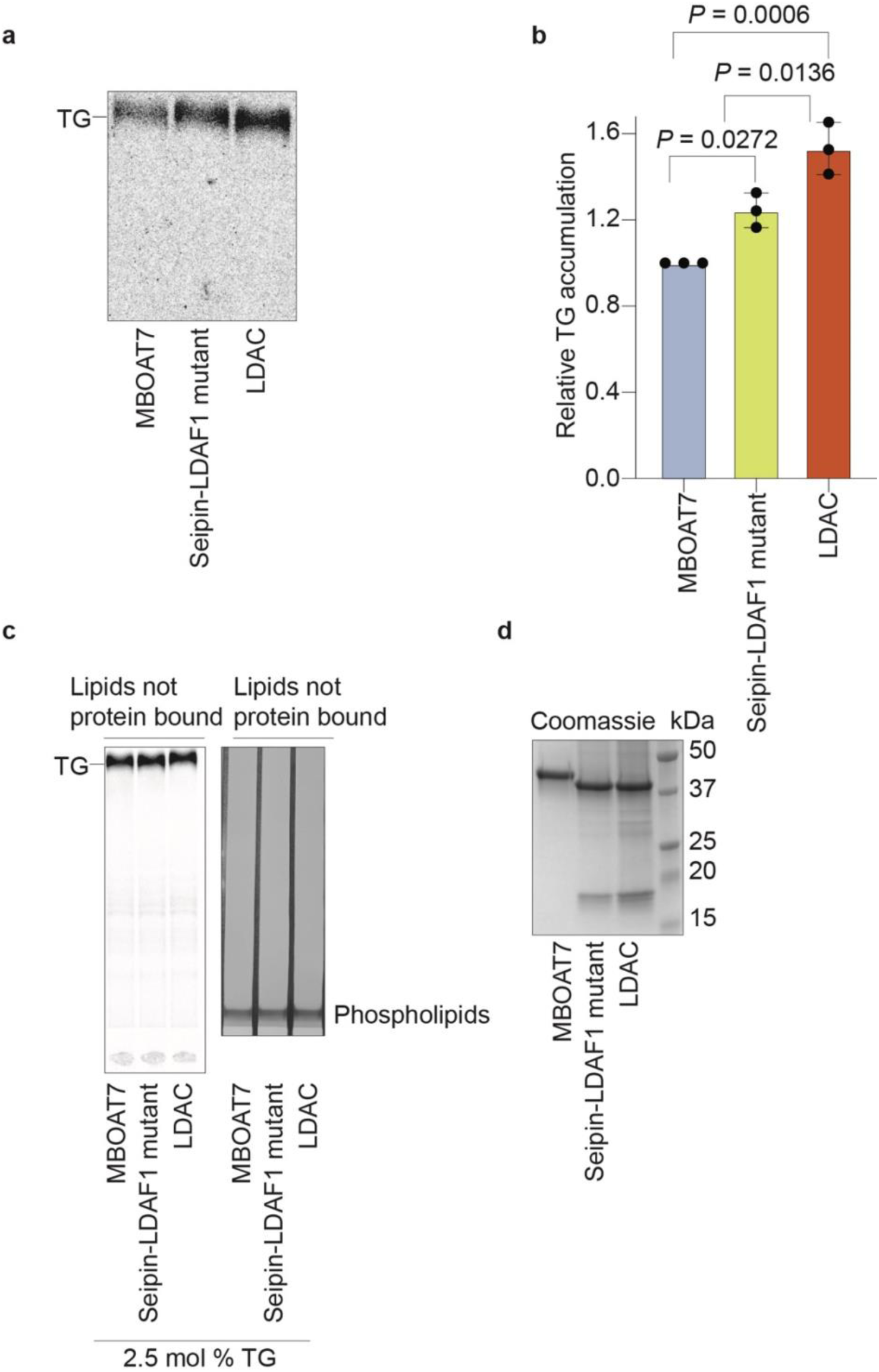
TG binding assay indicates that conserved residues in LDAF1 are important for TG interaction. **a,** LDAC binds TG, and the LDAC mutant has reduced affinity for TG. TG binding assay with purified proteins at 2.5 mol % TG. Elution of re-purified proteins was loaded on TLC to separate radiolabeled TG. **b**, Quantitation of signal intensity corresponding to TG in TLCs. One-way ANOVA was performed (mean ± s.d., n=3). **c,** All samples have equal amounts of phospholipids and TG as starting material. Flow-through of re-purified proteins from a, loaded on TLC. Radiolabeled TG was detected, as well as phospholipids that were stained with iodine vapor. **d,** Equal amount of proteins was used in each condition. Coomassie-stained SDS-page gel of purified proteins used in a.

**Extended Data Fig. 6.**
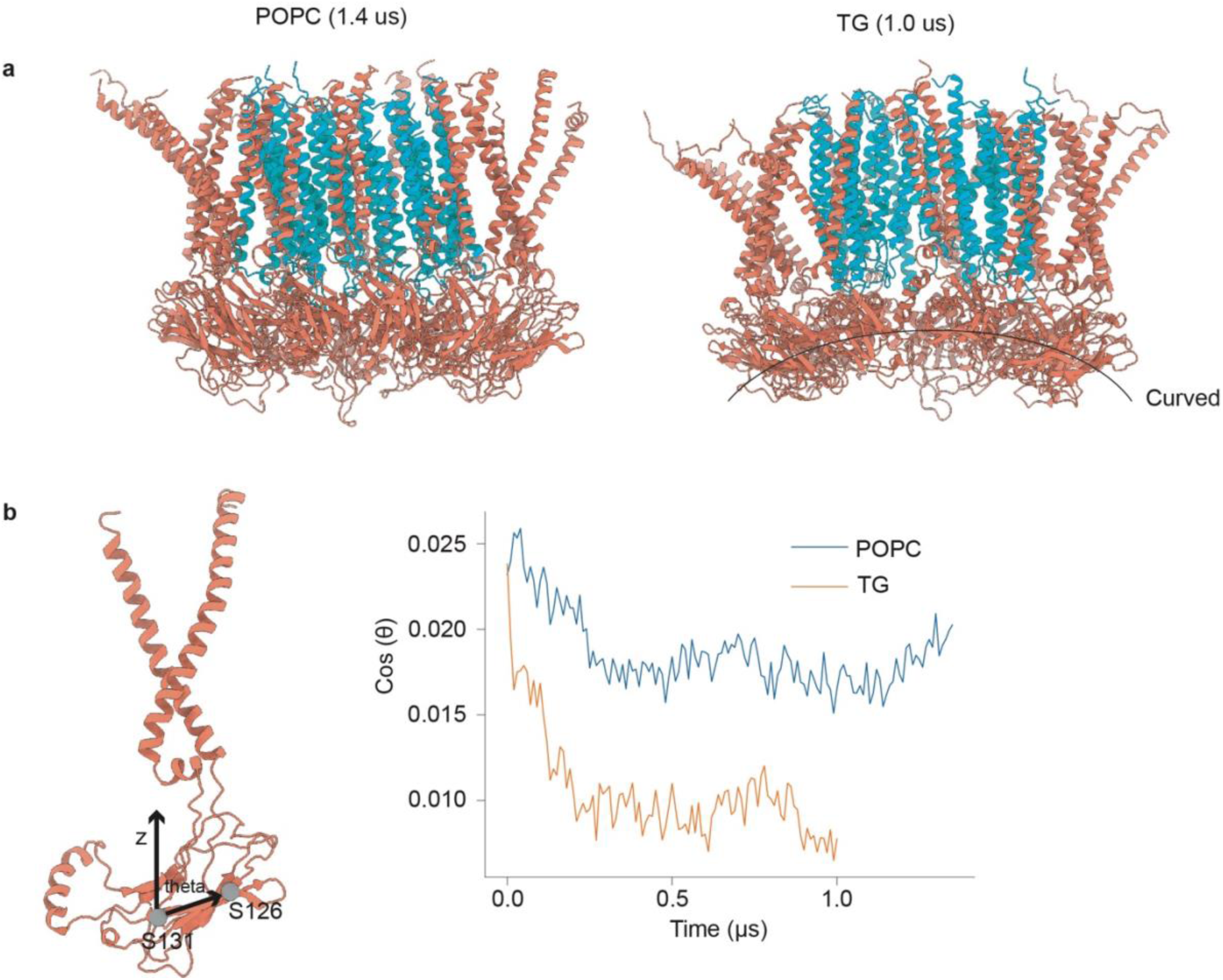
Phase-separated TG within the LDAC alters the orientation of the lumenal domain of seipin. **a**, LDAC conformations at 1.4 µs without nucleated TG (left) and at 1.0 µs with nucleated TG present (right). **b**, Angle between the membrane normal and the vector connecting residues Ser126 and Ser131.

**Extended Data Fig. 7.**
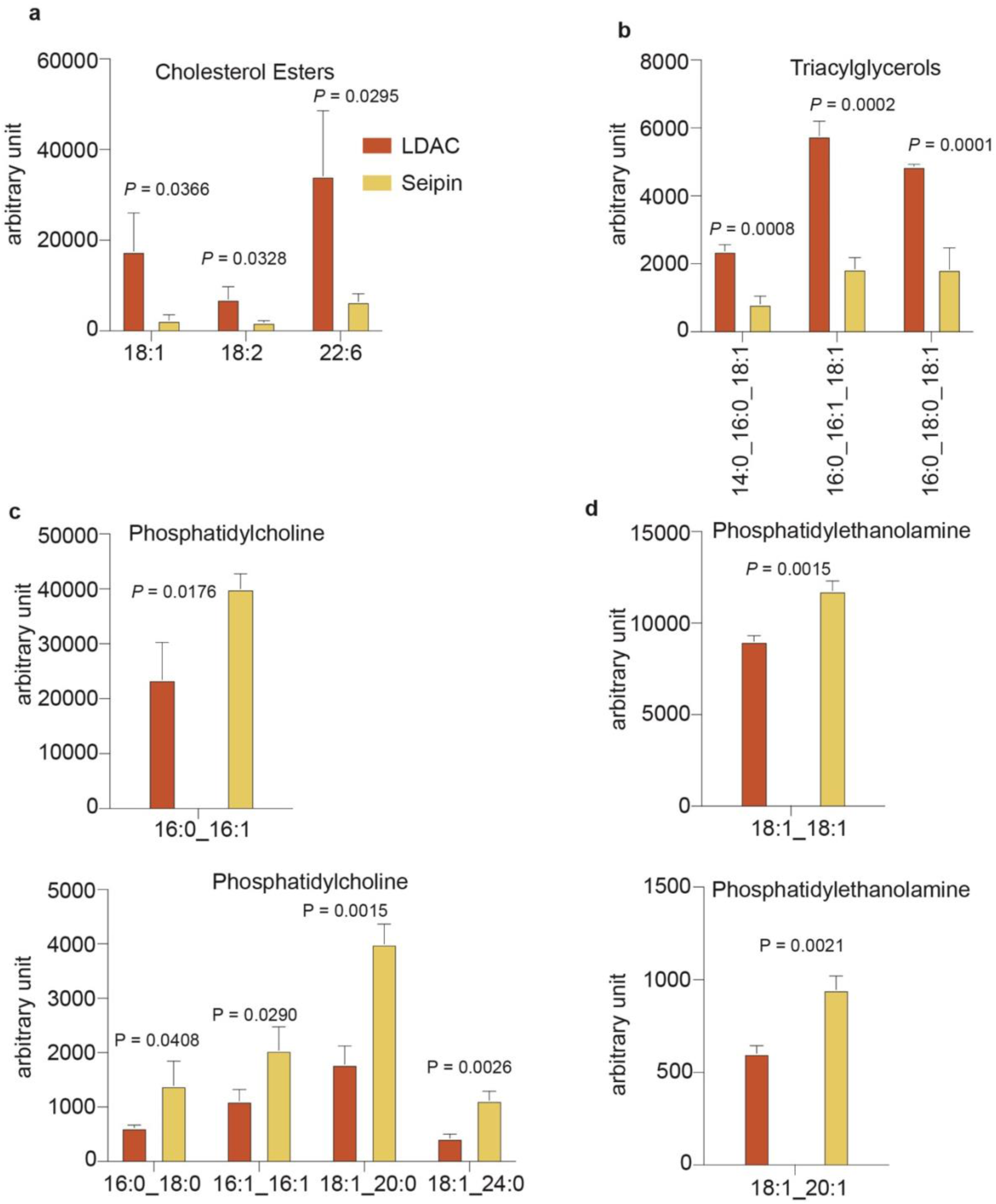
Lipidomic analysis indicates that the LDAC limits phospholipids while allowing TG accumulation. **a,** Cholesterol esters were enriched in LDAC, compared to seipin alone. Cholesterol esters abundance in isolated protein sample. Multiple t-test (mean ± s.d., n=3). **b,** Triacylglycerols are enriched in LDAC, compared to seipin alone. Triacylglycerols abundance in isolated protein samples. Multiple t-test (mean ± s.d., n=3). **c,** Phosphatidylcholine is enriched in seipin alone, compared to LDAC. Phosphatidylcholine abundance in isolated protein sample. Multiple t-test (mean ± s.d., n=3). **d,** Phosphatidylethanolamine is enriched in seipin alone compared to LDAC. Phosphatidylethanolamine abundance in isolated protein sample. Multiple t-test (mean ± s.d., n=3).

**Extended Data Fig. 8.**
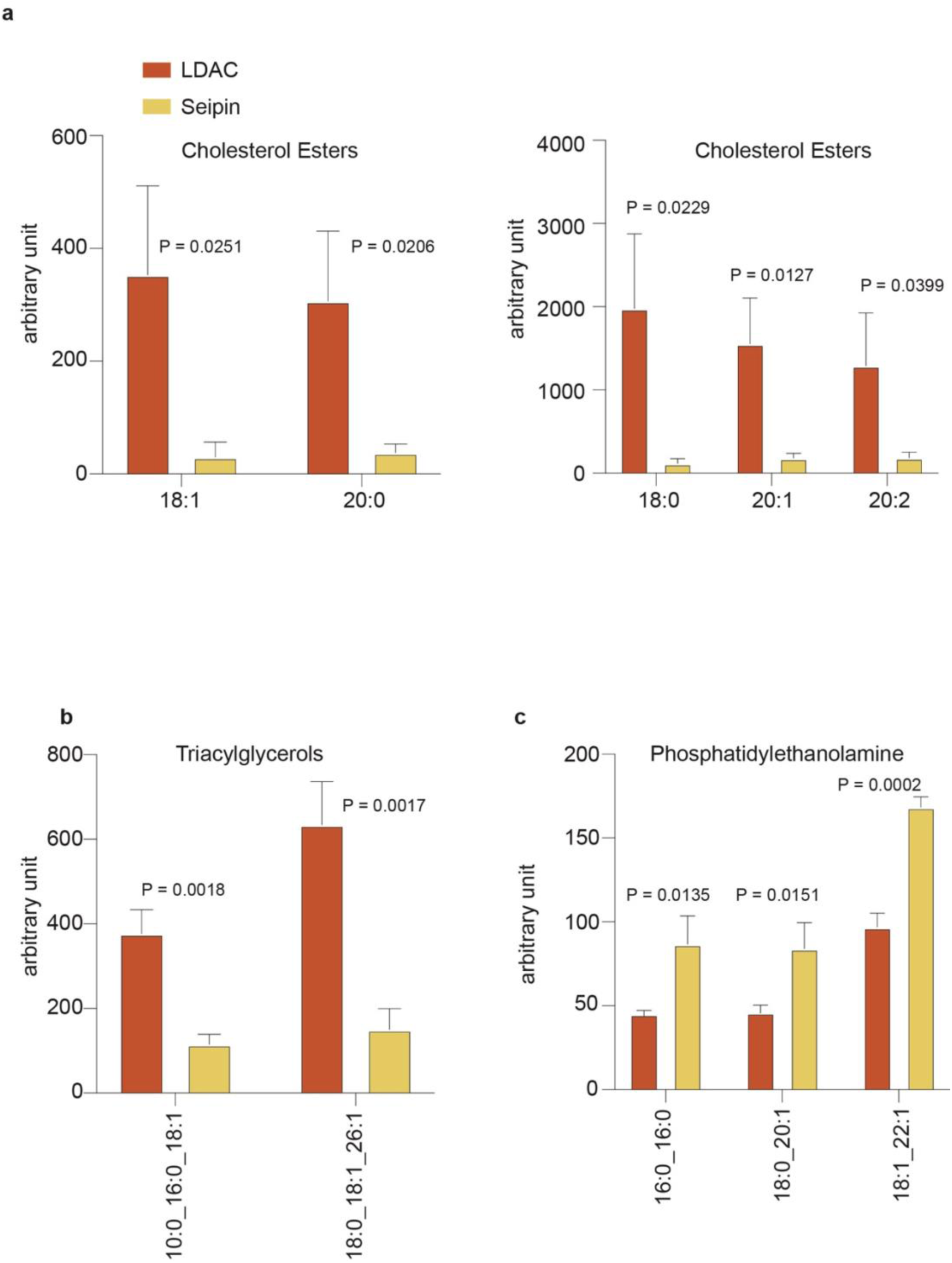
Lipidomic analyses indicate that the LDAC limits phospholipid access while allowing TG accumulation. **a**, Cholesterol esters are enriched in LDAC, compared to seipin alone. Cholesterol esters abundance in isolated protein sample. Multiple t-test (mean ± s.d., n=3). **b,** Triacylglycerols are enriched in LDAC, compared to seipin alone. Triacylglycerols abundance in isolated protein samples. Multiple t-test (mean ± s.d., n=3). **c,** Phosphatidylethanolamine is enriched in seipin alone, compared to LDAC. Phosphatidylethanolamine abundance in isolated protein sample. Multiple t-test (mean ± s.d., n=3).

## Acknowledgments

We thank members of the Farese & Walther laboratory and specifically Drs. Henning Arlt and Jeeyun Chung for helpful discussions, Dr. Ritchie Ly for critically reading the manuscript, Drs. Chandramohan Chitraju and Wei-Chung Tang for experimental consultation, and M. J. de la Cruz of the Structural Biology Core Facility at Memorial Sloan Kettering Cancer Center and the New York Structural Biology Center for help with cryo-EM data acquisition. We thank the high-performance computer center at Memorial Sloan Kettering Cancer Center for providing computational resources for cryo-EM and MD simulations. We thank Gary Howard for editorial assistance. We acknowledge NIH/NCI Cancer Center Support Grant (P30 CA008748) to MSKCC. This work was supported by National Institute of Health grant R01GM124348 (to R.V.F.) and R01GM063796 (to S.K. and G.A.V.). Yohannes Ambaw was supported by postdoctoral fellowship from the Bluefield Project. Tobias C. Walther is a Howard Hughes Medical Institute Investigator.

## Author contributions

P.C.M., R.V.F., and T.C.W. designed and supervised the project. P.C.M. carried out and analyzed experiments. S.K. performed MD simulations. S.K. and G.A.V. analyzed MD simulations. Y.A. performed lipidomic experiments and data analysis. P.C.M. and S.K. created figures. P.C.M., R.V.F., and T.C.W. wrote the original manuscript draft. P.C.M., S.K., R.V.F., and T.C.W. edited the manuscript, which all authors read.

## Competing interests declaration

The authors declare no competing interests.

## Additional information

Correspondence and requests for materials should be addressed to Robert V. Farese Jr. and Tobias C. Walther.

Reprints and permissions information is available at www.nature.com/reprints.

## Extended figure/table legends

**Table 1.**
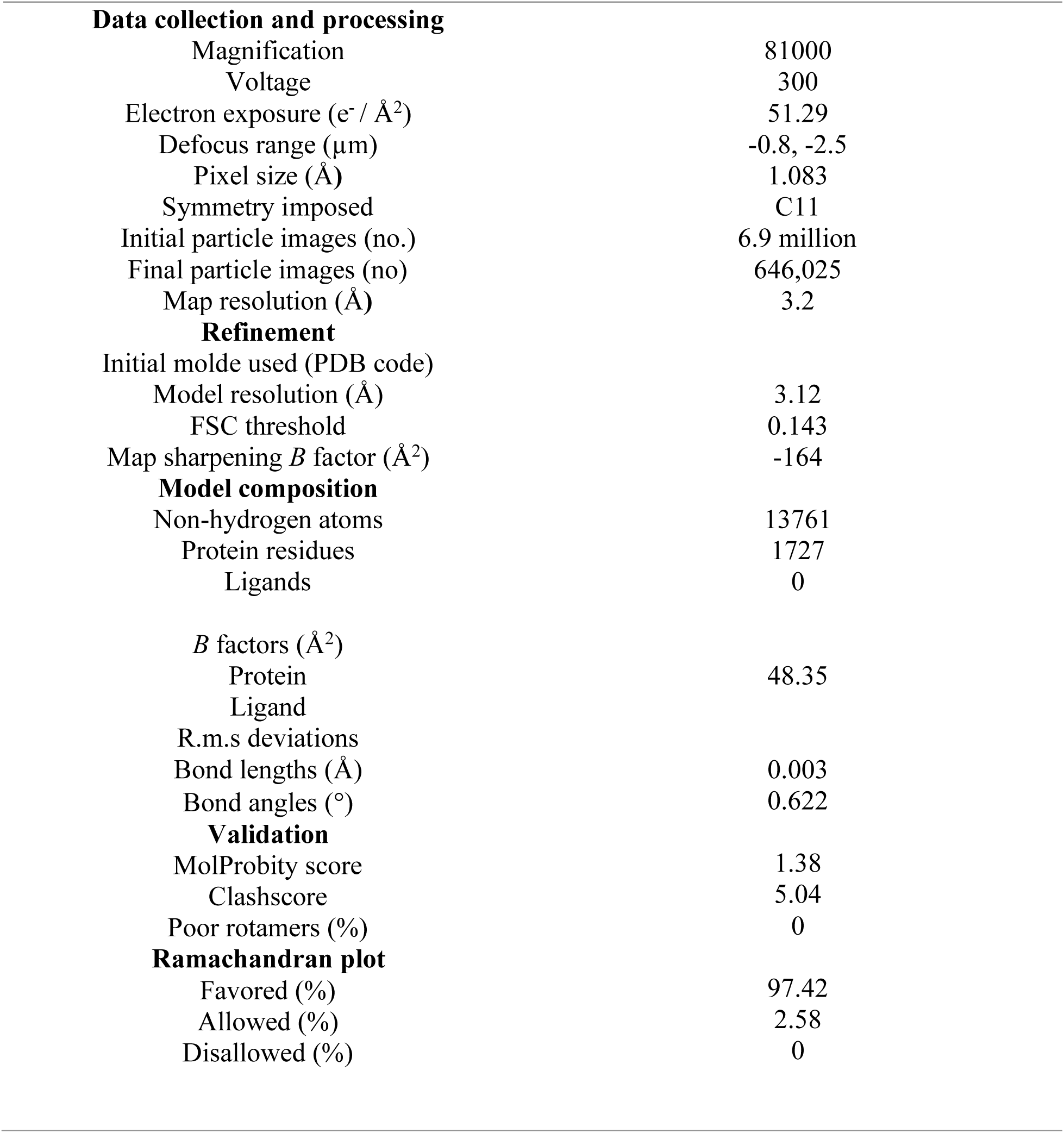
Cryo-EM data collection, refinement and validation statistics.

**Table 2.** Simulation setup.

| **System** | **Resolution** | **TG mol%** | **POPC:TG in upper leaflet** | **POPC:TG in lower leaflet** | **Simulation length** |
| --- | --- | --- | --- | --- | --- |
| LDAF1 protomer | CG | 0.25 mol% | 200:1 | 200:0 | 20 μs |
| Seipin | CG | 5 mol% | 3800:200 | 3725:200 | 10 μs |
| Seipin | CG | 2 mol% | 4000:80 | 3920:80 | 10 μs |
| Seipin | CG | 1 mol% | 4000:40 | 3920:40 | 10 μs |
| Seipin + LDAF1 | CG | 5 mol% | 3800:200 | 3800:200 | 10 μs |
| Seipin + LDAF1 | CG | 2 mol% | 3920:80 | 3920:80 | 10 μs |
| Seipin + LDAF1 | CG | 1 mol% | 3960:40 | 3960:40 | 10 μs |
| Seipin + LDAF1 | CG | 5 mol% | 950:50 | 950:50 | 10 μs |
| Seipin + LDAF1 | AA | 5 mol% | 950:50 | 950:50 | 1 μs |
| Seipin + LDAF1 | AA | 0 mol% | 700:0 | 700:0 | 1.4 μs |

## Notes

### Competing Interest Statement

The authors have declared no competing interest.

## References

1 Castro, I. G. et al. Promethin Is a Conserved Seipin Partner Protein. Cells 8 (2019). 10.3390/cells8030268

2 Chung, J. et al. LDAF1 and Seipin Form a Lipid Droplet Assembly Complex. Dev Cell 51, 551–563 e557 (2019). 10.1016/j.devcel.2019.10.006

3 Rogers, M. A. et al. Acyl-CoA:cholesterol acyltransferases (ACATs/SOATs): Enzymes with multiple sterols as substrates and as activators. J Steroid Biochem Mol Biol 151, 102–107 (2015). 10.1016/j.jsbmb.2014.09.008

4 Cases, S. e. a. Identification of a gene encoding an acyl CoA:diacylglycerol acyltransferase, a key enzyme in triacylglycerol synthesis. Proc Natl Acad Sci U S A 95, 13018–13023 (1998). 10.1073/pnas.95.22.13018

5 Cases, S. et al. Cloning of DGAT2, a second mammalian diacylglycerol acyltransferase, and related family members. J Biol Chem 276, 38870–38876 (2001). 10.1074/jbc.M106219200

6 Sui, X. et al. Structure and catalytic mechanism of a human triacylglycerol-synthesis enzyme. Nature 581, 323–328 (2020). 10.1038/s41586-020-2289-6

7 Wang, L. et al. Structure and mechanism of human diacylglycerol O-acyltransferase 1. Nature 581, 329–332 (2020). 10.1038/s41586-020-2280-2

8 Small, J. A. H. a. D. M. Solubilization and localization of triolein in phosphatidylcholine bilayers: A 13C NMR study. Proc Natl Acad Sci U S A Vol. 78, 6878–6882 (1981). 10.1073/pnas.78.11.6878

9 Thiam, A. R. & Ikonen, E. Lipid Droplet Nucleation. Trends Cell Biol 31, 108–118 (2021). 10.1016/j.tcb.2020.11.006

10 Holtta-Vuori, M., Salo, V. T., Ohsaki, Y., Suster, M. L. & Ikonen, E. Alleviation of seipinopathy-related ER stress by triglyceride storage. Hum Mol Genet 22, 1157–1166 (2013). 10.1093/hmg/dds523

11 Bi, J. et al. Seipin promotes adipose tissue fat storage through the ER Ca(2)(+)-ATPase SERCA. Cell Metab 19, 861–871 (2014). 10.1016/j.cmet.2014.03.028

12 Chitraju, C. et al. Triglyceride Synthesis by DGAT1 Protects Adipocytes from Lipid-Induced ER Stress during Lipolysis. Cell Metab 26, 407–418 e403 (2017). 10.1016/j.cmet.2017.07.012

13 Cui, X. et al. Seipin ablation in mice results in severe generalized lipodystrophy. Hum Mol Genet 20, 3022–3030 (2011). 10.1093/hmg/ddr205

14 Eisenberg-Bord, M. et al. Identification of seipin-linked factors that act as determinants of a lipid droplet subpopulation. J Cell Biol 217, 269–282 (2018). 10.1083/jcb.201704122

15 Teixeira, V. et al. Regulation of lipid droplets by metabolically controlled Ldo isoforms. J Cell Biol 217, 127–138 (2018). 10.1083/jcb.201704115

16 Magre, J. et al. Identification of the gene altered in Berardinelli-Seip congenital lipodystrophy on chromosome 11q13. Nat Genet 28, 365–370 (2001). 10.1038/ng585

17 Szymanski K. M. et al. The lipodystrophy protein seipin is found at endoplasmic reticulum lipid droplet junctions and is important for droplet morphology. Proc Natl Acad Sci U S A 104 (2007). 10.1073/pnas.0704154104

18 Salo, V. T. et al. Seipin regulates ER-lipid droplet contacts and cargo delivery. EMBO J 35, 2699–2716 (2016). 10.15252/embj.201695170

19 Sui, X. et al. Cryo-electron microscopy structure of the lipid droplet-formation protein seipin. J Cell Biol 217, 4080–4091 (2018). 10.1083/jcb.201809067

20 Yan, R. et al. Human SEIPIN Binds Anionic Phospholipids. Dev Cell 47, 248–256 e244 (2018). 10.1016/j.devcel.2018.09.010

21 Klug, Y. A. et al. Mechanism of lipid droplet formation by the yeast Sei1/Ldb16 Seipin complex. Nat Commun 12, 5892 (2021). 10.1038/s41467-021-26162-6

22 Arlt, H. et al. Seipin forms a flexible cage at lipid droplet formation sites. Nat Struct Mol Biol 29, 194–202 (2022). 10.1038/s41594-021-00718-y

23 Pagac, M. et al. SEIPIN Regulates Lipid Droplet Expansion and Adipocyte Development by Modulating the Activity of Glycerol-3-phosphate Acyltransferase. Cell Rep 17, 1546–1559 (2016). 10.1016/j.celrep.2016.10.037

24 Fei, W. et al. A role for phosphatidic acid in the formation of “supersized” lipid droplets. PLoS Genet 7, e1002201 (2011). 10.1371/journal.pgen.1002201

25 Sim, M. F. et al. The human lipodystrophy protein seipin is an ER membrane adaptor for the adipogenic PA phosphatase lipin 1. Mol Metab 2, 38–46 (2012). 10.1016/j.molmet.2012.11.002

26 Wolinski, H. et al. Seipin is involved in the regulation of phosphatidic acid metabolism at a subdomain of the nuclear envelope in yeast. Biochim Biophys Acta 1851, 1450–1464 (2015). 10.1016/j.bbalip.2015.08.003

27 Ding, L. et al. Seipin regulates lipid homeostasis by ensuring calcium-dependent mitochondrial metabolism. EMBO J 37 (2018). 10.15252/embj.201797572

28 Guyard, V. et al. ORP5 and ORP8 orchestrate lipid droplet biogenesis and maintenance at ER-mitochondria contact sites. J Cell Biol 221 (2022). 10.1083/jcb.202112107

29 Combot, Y. et al. Seipin localizes at endoplasmic-reticulum-mitochondria contact sites to control mitochondrial calcium import and metabolism in adipocytes. Cell Rep 38, 110213 (2022). 10.1016/j.celrep.2021.110213

30 Prasanna, X. et al. Seipin traps triacylglycerols to facilitate their nanoscale clustering in the endoplasmic reticulum membrane. PLoS Biol 19, e3000998 (2021). 10.1371/journal.pbio.3000998

31 Zoni, V. et al. Seipin accumulates and traps diacylglycerols and triglycerides in its ring-like structure. Proc Natl Acad Sci U S A 118 (2021). 10.1073/pnas.2017205118

32 Kim, S. et al. Seipin transmembrane segments critically function in triglyceride nucleation and lipid droplet budding from the membrane. Elife 11 (2022). 10.7554/eLife.75808

33 Abramson, J. et al. Accurate structure prediction of biomolecular interactions with AlphaFold 3. Nature 630, 493–500 (2024). 10.1038/s41586-024-07487-w

34 Marks, D. S., Hopf, T. A. & Sander, C. Protein structure prediction from sequence variation. Nat Biotechnol 30, 1072–1080 (2012). 10.1038/nbt.2419

35 Kim, S. Backmapping with Mapping and Isomeric Information. The Journal of Physical Chemistry B 127, 10488–10497 (2023). 10.1021/acs.jpcb.3c05593

36 Kim, S. et al. Seipin transmembrane segments critically function in triglyceride nucleation and lipid droplet budding from the membrane. eLife 11, e75808 (2022). 10.7554/eLife.75808

37 Salo, V. T. et al. Seipin Facilitates Triglyceride Flow to Lipid Droplet and Counteracts Droplet Ripening via Endoplasmic Reticulum Contact. Developmental Cell 50, 478–493.e479 (2019). 10.1016/j.devcel.2019.05.016

38 Renne, M. F., Corey, R. A., Ferreira, J. V., Stansfeld, P. J. & Carvalho, P. Seipin concentrates distinct neutral lipids via interactions with their acyl chain carboxyl esters. J Cell Biol 221 (2022). 10.1083/jcb.202112068

39 Shen, W. J. et al. Mice deficient in ER protein seipin have reduced adrenal cholesteryl ester lipid droplet formation and utilization. J Lipid Res 63, 100309 (2022). 10.1016/j.jlr.2022.100309

40 Wang, H. et al. Seipin is required for converting nascent to mature lipid droplets. Elife 5 (2016). 10.7554/eLife.16582

41 Choudhary, V., El Atab, O., Mizzon, G., Prinz, W. A. & Schneiter, R. Seipin and Nem1 establish discrete ER subdomains to initiate yeast lipid droplet biogenesis. J Cell Biol 219 (2020). 10.1083/jcb.201910177

42 Li, C. e. a. Adipogenin dictates adipose tissue expansion by facilitating the assembly of a dodecameric seipin complex. BioRxiv (2024). 10.1101/2024.07.25.605195

## Methods references

43 Dezi, M., Di Cicco, A., Bassereau, P. & Levy, D. Detergent-mediated incorporation of transmembrane proteins in giant unilamellar vesicles with controlled physiological contents. Proc Natl Acad Sci U S A 110, 7276–7281 (2013). 10.1073/pnas.1303857110

44 Angelova, M. I. e. a. Preparation of giant vesicles by external AC electric fields. Kinetics and applications. Progress in Colloid & Polymer Science 89, 127–131 (1992). 10.1007/BFb0116295

45 Schindelin, J., et al. Fiji: an open-source platform for biological-image analysis. Nat Methods 9, 676–682 (2012). 10.1038/nmeth.2019

46 Suloway, C. et al. Automated molecular microscopy: the new Leginon system. J Struct Biol 151, 41–60 (2005). 10.1016/j.jsb.2005.03.010

47 Punjani A. et al. cryoSPARC: algorithms for rapid unsupervised cryo-EM strcuture determination. Nature Methods 14, 290–296 (2017). 10.1038/nmeth.4169nature

48 Bepler, T. et al. Positive-unlabeled convolutional neural networks for particle picking in cryo-electron micrographs. Nat Methods 16, 1153–1160 (2019). 10.1038/s41592-019-0575-8

49 Pettersen, E. F. et al. UCSF ChimeraX: Structure visualization for researchers, educators, and developers. Protein Sci 30, 70–82 (2021). 10.1002/pro.3943

50 Emsley, P. & Cowtan, K. Coot: model-building tools for molecular graphics. Acta Crystallogr D Biol Crystallogr 60, 2126–2132 (2004). 10.1107/S0907444904019158

51 Adams, P. D. e. a. PHENIX: building new software for automated crystallographic strcuture determination. Acta Crystallogr D Biol Crystallogr 58, 1948–1954 (2002). 10.1107/s0907444902016657

52 Williams, C. J. et al. MolProbity: More and better reference data for improved all-atom structure validation. Protein Sci 27, 293–315 (2018). 10.1002/pro.3330

53 Hamed, A. et al. LipidCruncher: An open-source web application for processing, visualizing, and analyzinglipidomic data. BioRxiv (2025). 10.1101/2025.04.28.650893

54 Scheich, C., Kummel, D., Soumailakakis, D., Heinemann, U. & Bussow, K. Vectors for co-expression of an unrestricted number of proteins. Nucleic Acids Res 35, e43 (2007). 10.1093/nar/gkm067

55 Abraham, M. J. et al. GROMACS: High performance molecular simulations through multi-level parallelism from laptops to supercomputers. SoftwareX 1-2, 19–25 (2015). 10.1016/j.softx.2015.06.001

56 Van Der Spoel, D. e. a. GROMACS: Fast, flexible, and free. Journal of Computational Chemistry 26, 1701–1718 (2005). 10.1002/jcc.20291

57 Huang, J. et al. CHARMM36m: an improved force field for folded and intrinsically disordered proteins. Nature Methods 14, 71–73 (2017). 10.1038/nmeth.4067

58 Klauda, J. B. et al. Update of the CHARMM All-Atom Additive Force Field for Lipids: Validation on Six Lipid Types. The Journal of Physical Chemistry B 114, 7830–7843 (2010). 10.1021/jp101759q

59 de Jong, D. H. et al. Improved Parameters for the Martini Coarse-Grained Protein Force Field. J Chem Theory Comput 9, 687–697 (2013). 10.1021/ct300646g

60 Marrink, S. J., Risselada, H. J., Yefimov, S., Tieleman, D. P. & de Vries, A. H. The MARTINI Force Field: Coarse Grained Model for Biomolecular Simulations. The Journal of Physical Chemistry B 111, 7812–7824 (2007). 10.1021/jp071097f

61 Monticelli, L. e. a. The MARTINI Coarse-Grained Force Field: Extension to Proteins. Journal of Chemival Theory and Computation 4, 819–834 (2008). 10.1021/ct700324x

62 Campomanes, P., Prabhu, J., Zoni, V. & Vanni, S. Recharging your fats: CHARMM36 parameters for neutral lipids triacylglycerol and diacylglycerol. Biophysical Reports 1 (2021). 10.1016/j.bpr.2021.100034

63 Khandelia, H., Duelund, L., Pakkanen, K. I. & Ipsen, J. H. Triglyceride blisters in lipid bilayers: implications for lipid droplet biogenesis and the mobile lipid signal in cancer cell membranes. PLoS One 5, e12811 (2010). 10.1371/journal.pone.0012811

64 Nosé, S. A unified formulation of the constant temperature molecular dynamics methods. The Journal of Chemical Physics 81, 511–519 (1984). 10.1063/1.447334

65 Hoover, W. G. Canonical dynamics: Equilibrium phase-space distributions. Physical Review A 31, 1695–1697 (1985). 10.1103/PhysRevA.31.1695

66 Parrinello, M. & Rahman, A. Polymorphic transitions in single crystals: A new molecular dynamics method. Journal of Applied Physics 52, 7182–7190 (1981). 10.1063/1.328693

67 Bernetti, M. B., G. Pressure control using stochastic cell rescaling. The Journal of Chemical Physics 153 (2020). 10.1063/5.0020514

68 Bussi, G., Donadio, D. & Parrinello, M. Canonical sampling through velocity rescaling. The Journal of Chemical Physics 126 (2007). 10.1063/1.2408420

69 Kim, S. Backmapping with Mapping and Isomeric Information. J Phys Chem B 127, 10488–10497 (2023). 10.1021/acs.jpcb.3c05593

70 Kim, S. All-Atom Membrane Builder via Multiscale Simulation. J Chem Inf Model 64, 7077–7085 (2024). 10.1021/acs.jcim.4c01059

71 Michaud-Agrawal, N., Denning, E. J., Woolf, T. B. & Beckstein, O. MDAnalysis: a toolkit for the analysis of molecular dynamics simulations. J Comput Chem 32, 2319–2327 (2011). 10.1002/jcc.21787

